# Conserved degronome features governing quality control associated proteolysis

**DOI:** 10.1101/2022.04.06.487275

**Authors:** Bayan Mashahreh, Shir Armony, Kristoffer Enøe Johansson, Alon Chappleboim, Nir Friedman, Richard G. Gardner, Rasmus Hartmann-Petersen, Kresten Lindorff-Larsen, Tommer Ravid

## Abstract

The eukaryotic proteome undergoes constant surveillance by quality control systems that either sequester, refold, or eliminate aberrant proteins by ubiquitin-dependent mechanisms. Ubiquitin-conjugation necessitates the recognition of degradation determinants, termed degrons, by their cognate E3 ubiquitin-protein ligases. To learn about the distinctive properties of quality control degrons, we performed an unbiased peptidome stability screen in yeast. The search identified a large cohort of proteome-derived degrons, some of which exhibited broad E3 ligase specificity. Consequent application of a machine-learning algorithm established constraints governing degron potency, including the amino acid composition and secondary structure propensities. According to the set criteria, degrons with transmembrane domain-like characteristics are the most probable sequences to act as degrons. Similar quality control degrons were identified in viral and human proteins, suggesting conserved degradation mechanisms. Altogether, the emerging data indicate that transmembrane domain-like degron features have been preserved in evolution as key quality control determinants of proteins’ half-life.

## Introduction

Intracellular protein quality control (PQC) is a principal regulatory mechanism for the maintenance of protein homeostasis ^1^. PQC systems continuously survey the proteome and execute a triage of unfolded protein states, the result of which is either refolding or, if beyond repair, sequestration or degradation of aberrant proteins ^2,3^. Protein refolding and sequestration are primarily mediated by molecular chaperones ^4–6^, while the Ubiquitin-Proteasome System (UPS) executes quality control-associated proteolysis (QCAP) ^7^.

A key to understanding protein homeostasis is deciphering the mode by which QCAP pathways discern the folding state of proteins. It has been established that the ubiquitin conjugation system, via E3 ubiquitin-protein ligases and auxiliary chaperones, recognize degradation determinants termed degrons ^8^ that constitute inherent sequences and structural features, as well as acquired post-translational modifications ^9^. To date, degrons have been mostly identified through studies of regulated protein degradation mechanisms, such as those involved in cell division and differentiation, as well as in many cancer-related diseases ^10–12^. These studies identified inherent degrons as short motifs, such as the destruction box of cyclins, as well as acquired degrons that are activated by phosphorylation or other post-translational modifications. However, the repertoire of known degrons cannot explain the large diversity in half-lives exhibited by the proteome ^13,14^.

Our earlier work exposed the large sequence heterogenicity of the cellular degron landscape (degronome) in yeast, which led to the proposition that the majority of the eukaryotic proteome contains cryptic QCAP degrons that may become exposed naturally or under misfolding conditions, such as cellular and environmental stresses ^15,16^. These degrons target protein ubiquitination via the activity of a relatively small number of designated QCAP E3 ligases, suggesting that each recognizes a large and possibly diverse set of substrates ^17^. Furthermore, QCAP E3 enzymes can act redundantly in the ubiquitination of their substrates, seemingly exhibiting overlapping recognition mechanisms ^18–20^. However, the significance of this functional redundancy is not yet fully understood.

Here we describe a yeast-adapted Global Protein Stability (GPS)-peptidome technology (yGPS-P) originally established for the discovery of degrons in mammalian cell lines ^21–23^. By employing a peptide library fused to yGPS-P, we have identified multiple degron sequences that were subsequently analyzed using a machine learning algorithm. The resulting computer program termed Quality Control Degron Prediction (QCDPred)^24^ revealed amino acid preferences in QCAP degrons. The determined degron features were highly dependent on the overall hydrophobicity, and consistently transmembrane domains (TMDs) exhibit extreme degron potency, signifying their critical role in the degradation of integral membrane proteins prevented from entering the secretory pathway.

## Results

### Yeast-based GPS-peptidome technology

To set up a comprehensive degron discovery platform in the yeast *Saccharomyces cerevisiae* proteome, we applied a fluorescence-based GPS technology ^21^, previously developed in cultured human cells ^22,23^. yGPS-P utilizes a bicistronic gene expression system in which codon-optimized versions of yeast-enhanced Cherry (yeC) and yeast-enhanced GFP (yeG) are expressed from a single transcript. The two proteins are, however, translated separately due to the presence of an Internal Ribosome Entry Site (IRES) upstream to GFP that allows translation initiation in a cap-independent manner ^25^ (Figure 1a). A yeast GPS peptidome library (yGPS-P_lib_) is generated by subsequent in-frame insertion of proteome-derived DNA fragments downstream to GFP in a yGPS-P vector (Figure 1a). The plasmid library is transformed into yeast, followed by quantitative flow cytometry or Fluorescence-Activated Cell Sorting (FACS) (Figure 1b). As both Cherry and GFP are expressed from a single transcript yet translated independently, Cherry levels reflect the basal expression of the reporters while the ratio between GFP to Cherry (shown henceforth as yeG/yeC) determines the relative GFP protein level that is governed by the fused peptide.

**Figure 1.**
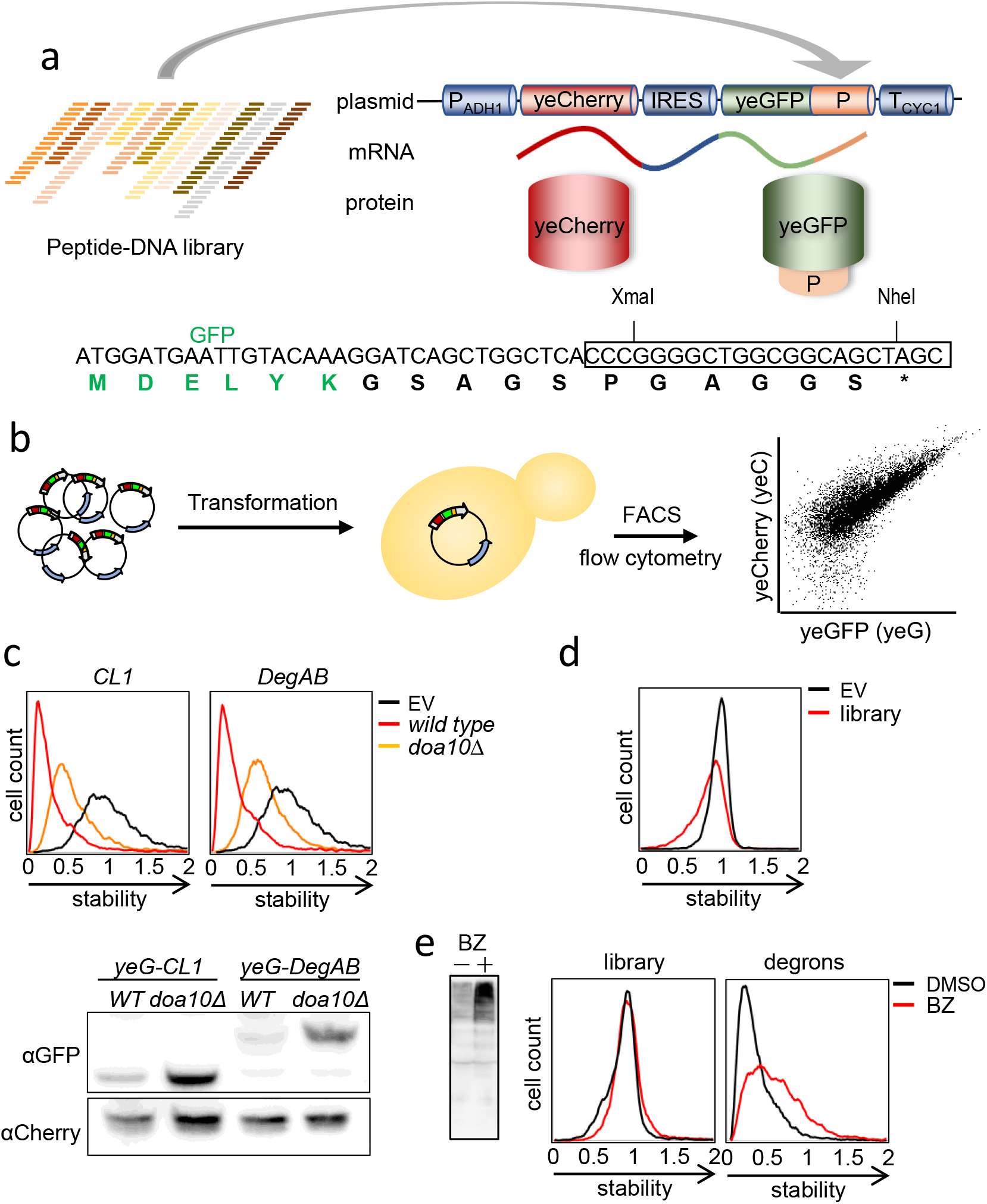
Principles and validation of the yGPS-peptidome method. (a) *Upper panel:* Schematic representation of yGPS-peptide library (yGPS-P_lib_) where a peptide-DNA library comprised of 51mer tiled DNA fragments from 335 proteins, represented by different colors, are cloned in-frame downstream to GFP in the yGPS-P vector. The single mRNA product consists of yeCherry-IRES-yeGFP-peptide. *Lower panel*: DNA and protein sequences illustrating the cloning site that pursues GFP and the pentameric linker. (b) Flowchart of the degron screen. yGPS-P_lib_ is transformed into yeast, followed by flow cytometry or FACS. (c) *Upper panel*: Flow cytometry histograms of *CL1* or *DegAB* degrons appended to yGPS-P, in *wild type* and *doa10∆* cells. Fluorescence emissions of 10,000 cells were determined for each condition. Scale: Median value of the yeG/yeC ratio in empty vector (EV) control was set as one. All other histograms were distributed accordingly. *Lower panel*: Immunoblot analysis of the levels of *CL1-*and *DegAB*-appended GFP compared to Cherry loading control. yeG: yeGFP, yeC: yeCherry. (d) Flow cytometry histogram of normalized yeG/yeC in yGPS-P_lib_ compared to empty vector control. Scale was set as shown in Figure 1C, *upper panel*. (e) Proteasome dependence of GFP-appended peptides. Cells expressing yGPS-P_lib_ were treated with 10µM Bortezomib (BZ) for four hours or with DMSO vehicle control. Cells were subjected to immunoblotting with anti-ubiquitin Abs (left panel), or to flow cytometry analysis (middle panel). Shown in the right panel is the effect of BZ on a pre-selected degron library yGPS-P_deg_ composed of top10% degrons. Scale was set as shown in Figure 1C, *upper panel*.

To validate the use of yGPS-P as a degron discovery platform, we examined the competence of two well-characterized QCAP degrons of the Doa10 E3 ligase, *CL1* and *DegAB* ^26–29^ to trigger degradation of the otherwise stable GFP. Doa10 is an endoplasmic reticulum (ER)-embedded enzyme that operates in ER-associated degradation (ERAD) ^30^. When the yeG/yeC ratio in cells expressing the fused degrons was compared to empty vector control, a more than 5-fold decrease was observed, presumably due to UPS-mediated proteolysis (Figure 1c, *upper panel*). An increase in yeG/yeC in *doa10∆* cells confirmed this assertion. That the increase in yeG/yeC is indeed a result of elevated GFP protein levels was demonstrated by immunoblot analysis of the corresponding fusion proteins (Figure 1c, *lower panel*).

As a source for a peptide library, we chose a subset of proteins, all components of multimeric protein complexes that potentially undergo QCAP triage ^32,33^. In total, 326 yeast proteins that are part of 23 different complexes were selected (Supplementary Data 1). These proteins operate in distinct cell compartments and the composition of amino acids and secondary structure elements of the selected proteins are similar to those of the entire yeast proteome (Supplementary Figures 1a, 1b). Consequently, 51-mer DNA fragments with 36-mer DNA overlaps (corresponding to 17 amino acid-length tiled peptides with 12 amino acid overlaps) were synthesized to give rise to a yGPS-P_lib_ (library) containing approximately 29,500 DNA fragments. yGPS-P_lib_ was transformed into yeast, followed by flow cytometry determination of the yeG/yeC ratio. The observed decrease in yeG/yeC in yGPS-P_lib_ (Figure 1d) indicates the presence of a destabilizing peptide population within the tested peptidome. To assess the contribution of degrons of the UPS to GFP destabilization, the effect of the reversible proteasome inhibitor Bortezomib (BZ) on yeG/yeC was determined. To this end, yGPS-P_lib_ was transformed into cells lacking the multidrug transporter *PDR5 (pdr5∆*) to increase drug sensitivity ^34^. Comparing mock- and drug-treated cells, we observed an increase in the overall cellular levels of ubiquitin conjugates (confirming proteasome inhibition), as well as a mild increase in yeG/yeC (Figure 1e, *left & middle panels*). A larger increase of yeG/yeC was observed when a pre-sorted top 10% degron-enriched population was tested (Fig. 1e, *right panel*, and Supplementary Figure 1c), confirming that changes in yeG/yeC accurately reflect susceptibility to UPS-mediated degradation.

### Mapping QCAP degrons using a machine-learning-based approach

To classify degron sequences within yGPS-P_lib_, mid-log-phase cells expressing the appended peptidome were separated by FACS into four gates according to yeG/yeC, each containing an equal cell number, and the identity and amounts of the peptides’ DNA in the different gates were determined by Next-Generation Sequencing (NGS) (Figure 2a). After filtering ambiguous peptides DNA from the NGS data, the contribution of 23,600 peptides to GFP stability was calculated based on their abundance in the different gates and each was assigned a Protein Stability Index (PSI) score ^21^ (Figure 2a and Supplementary Data 2). Overall, 9.5% of the analyzed peptidome had PSI values < 1.7 (on a scale of 1 – 4), suggesting a degron function (Figure 2a).

**Figure 2.**
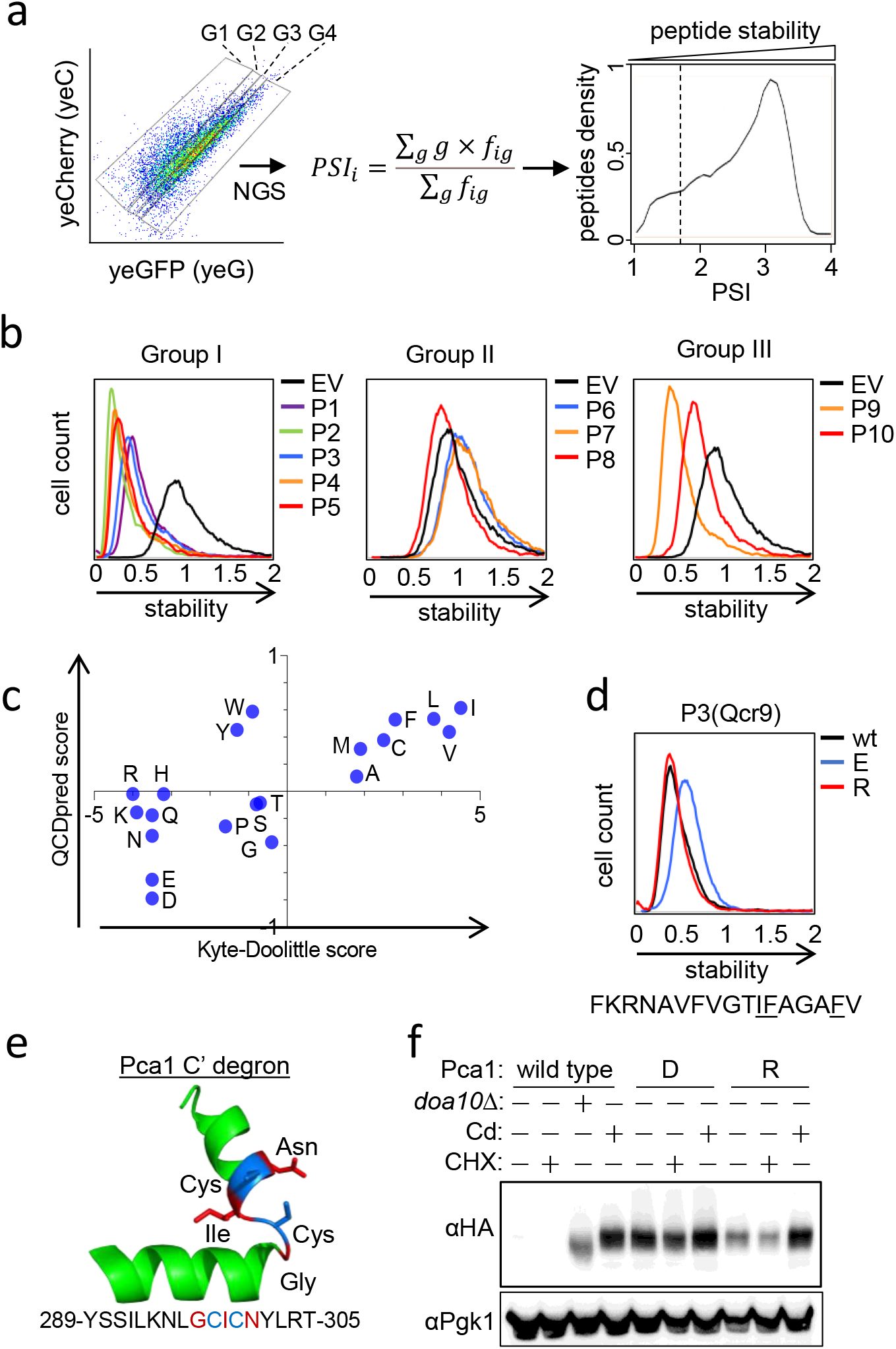
QCAP degrons functionality is largely defined by their amino acid composition. (a) Fluorescence-activated cell sorting of yGPS-P_lib_ into four gates (G1-4), each composed of 2.5 million cells, was followed by NGS and protein stability index (PSI) scoring of each peptide, leading to the formation of a peptide density map. Shown is a FACS illustration and gating of 10,000 cells. Degron cutoff at PSI < 1.7 is marked by a dashed line. (b) Validation of QCDPred-based degron predictions. Ten library peptides were re-cloned into yGPS-P, followed by flow cytometry analysis. The peptides were divided into three different groups: Group I-PSI ≤ 1.62, P ≥ 0.85; Group II-PSI > 1.62, P < 0.85; Group III-PSI ≤ 1.62, P < 0.85. Scale: Median value of the yeG/yeC ratio in empty vector (EV) control was set as one. All other histograms were distributed accordingly.(c) Scatter plot comparing Kyle-Doolittle hydrophobicity amino acid scores and QCDPred probabilities. (d) Flow cytometry histogram of intact or mutant P3 peptide in which three amino acids were replaced with either glutamate (E) or arginine (R) residues. Scale was set as shown in Figure 2B. (e) 3D structure of QCDPred-calculated cytosolic Pca1 degron (amino acids 289-305), based on AlphaFold Pca1 structural model #AF-p38360-F1 (Conf. (90 > pLDDT > 70). Marked by a blue color are the cysteine residues. Marked by a red color are the three amino acids that have been replaced with aspartate or arginine. pLDDT: per-residue confidence score on a scale of 0-100. (f) Immunoblot analysis of Pca1 protein levels. *Wild type* or *doa10∆* (*d∆*) cells, expressing the indicated HA-tagged Pca1 proteins were left intact or treated with 50 μM CdCl_2_ for 1 hour. To determine degradation dynamics, where indicated, the translation inhibitor cycloheximide (CHX) was added to cells for 15 min before cell harvesting. Pca1 levels were measured by immunoblotting using anti-HA Abs while Pgk1 levels served as a loading control. D: replacement of residues marked in 2E by red color with aspartate; R: replacement of residues marked in 2E by red color with Arginine.

To identify common motifs within the examined degrons, we next opted to employ a machine-learning algorithm, by using the PSI data to train a logistic regression model. The resulting computer program, termed QCDPred, provided each amino acid with a unique value, based on its predicted contribution to degron potency ^24^. Consequently, protein stability maps were formed for all proteins comprising the tested library that attained 100% coverage (N = 306), each includes experimental PSI values for each peptide, average PSI values for each amino acid, and QCDPred probability scores (Supplementary Figure 2).

To experimentally validate the high-throughput procedure and test the QCDPred model, we selected ten peptide sequences that we studied-one at a time -using the same reporter system and flow cytometry readout (Figure 2b, Table 1). Five peptides (P1–P5), which we predicted by QCDPred to have degron activity (group I), and three peptides that we predicted not to have degron activity (P6–P8; group II) were confirmed in this experiment. In contrast, two peptides (P9 and P10) were found to have some degron activity even though they were not predicted as degrons by QCDPred (group III). We also tested steady-state levels of selected peptides from each group and found a good correlation: Levels of group I and group III fusion peptides was significantly lower than that of group II (Supplementary Figure 3a). Moreover, treating the cells with a proteasome inhibitor (Bortezomib) restored GFP fluorescence for the degron sequences, demonstrating that the lowered fluorescence is due to proteasomal degradation, lending further support that QCDPred predicts proteasomal QCAP substrates (Supplementary Figure 3b).

**Table I.**
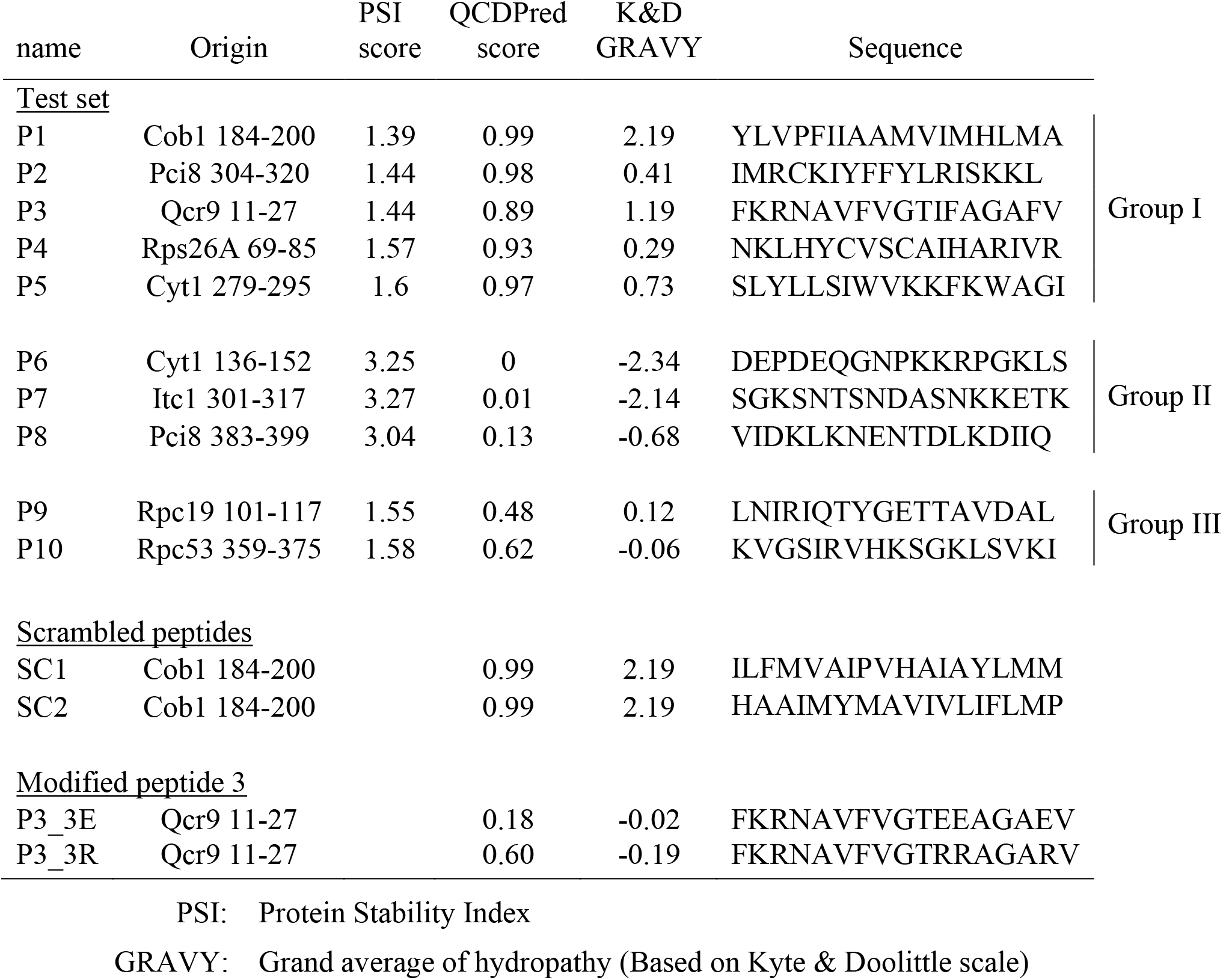
List of selected peptides, their origin, sequence, PSI and GRAVY scores

As projected from previous studies of QCAP degrons, QCDPred scores of most amino acid types are correlated with their Kyte-Doolittle hydrophobicity scores, and indeed, all hydrophobic amino acids contributed positively to QCAP degrons’ probability (Figure 2c). Conversely, QCDPred scores of the two negatively charged amino acids, glutamate and aspartate, were significantly lower than others, suggesting that the presence of negatively charged amino acids specifically interferes with a QCAP degron’s function. According to the prediction, inserting negatively charged amino acids into a peptide degron significantly reduces its QCDPred score (As an example, see Supplementary Table 1). This was confirmed experimentally by replacing three amino acids in a re-evaluated proteasome-dependent degron peptide P3 from the yeast protein Qcr9 with either glutamate or arginine (Figure 2d). We concluded that the classified QCAP degronome prefers hydrophobic residues while negatively charged amino acids are disfavored.

We next examined these assertions on a native QCAP degron from the P-type Cation-transporting ATPase Pca1, that was not in our tested degron cohort. Under standard growth conditions, Pca1 is constitutively degraded via Doa10, which recognizes a cysteine-enriched degron localized within amino acids 271-320 of the cytosolic and solvent-exposed N-terminal region of Pca1 ^35^. However, cadmium sensing by the degron enables Pca1 to circumvent ERAD ^35^. As this region is too long for precise analysis, we used QCDPred to locate a shorter sequence that defines the operational degron between amino acids 289-305, which was predicted by AlphaFold ^36,37^ to form an exposed helix-turn-helix structure (Figure 2e). Examining Pca1 steady-state levels, we observed that replacing three amino acids in the degron core (Figure 2e) with aspartate residues greatly stabilized the protein while replacing the same amino acids with arginine residues only showed moderate stabilization (Figure 2f). Importantly, mutant Pca1 still retained cadmium sensitivity (Figure 2f), indicating overall structural preservation. These data confirmed our assertion that negatively charged amino acids greatly interfere with native QCAP degron’s function. The data also demonstrated the capability of QCDPred to identify functional degrons within the proteome while in their physiological context.

### Transmembrane domains function as QCAP degrons

Intriguingly, besides the aforementioned cytosolic degron, QCDPred also assigned remarkably high degron probabilities (P ≥ 0.93) to Pca1 TMDs (Figure 3a). This was unexpected, not only because QCDPred was not programmed to consider protein topology, but also because TM proteins comprised only a minute proportion of the proteins included in the screen (1.79%) and hence their small contribution to the algorithm. The observation is, however, in line with PQC degrons being hydrophobic (Figure 2c). Thus, when applying QCDPred to the entire yeast proteome, the vast majority of TMDs were assigned as degrons (Figure 3b). Our experimental peptidome data agree with this prediction, demonstrating that 12 out of 13 peptides localized to TMDs of the inner mitochondrial cytochrome b-c1 respiratory complex ^38^, the only TM-embedded complex in our peptidome cohort (Supplementary Data 1), function as degrons (Figure 3c and Supplementary Figure 4). In line with these findings, most predicted TMD degrons in the yeast proteome were enriched in the highest QCDPred score range of 0.95-1.0 while the rest of the degrons were underrepresented in this range (Figure 3d). Thus, according to QCDPred, TMDs comprise the most potent QCAP degron sequences. Hence, the hydrophobic sequence and possibly structural resemblance to TMDs is likely a significant feature of QCAP degrons. TMDs themselves could be relevant as PQC degrons in cases when TM proteins fail to insert correctly in membranes (see next).

**Figure 3.**
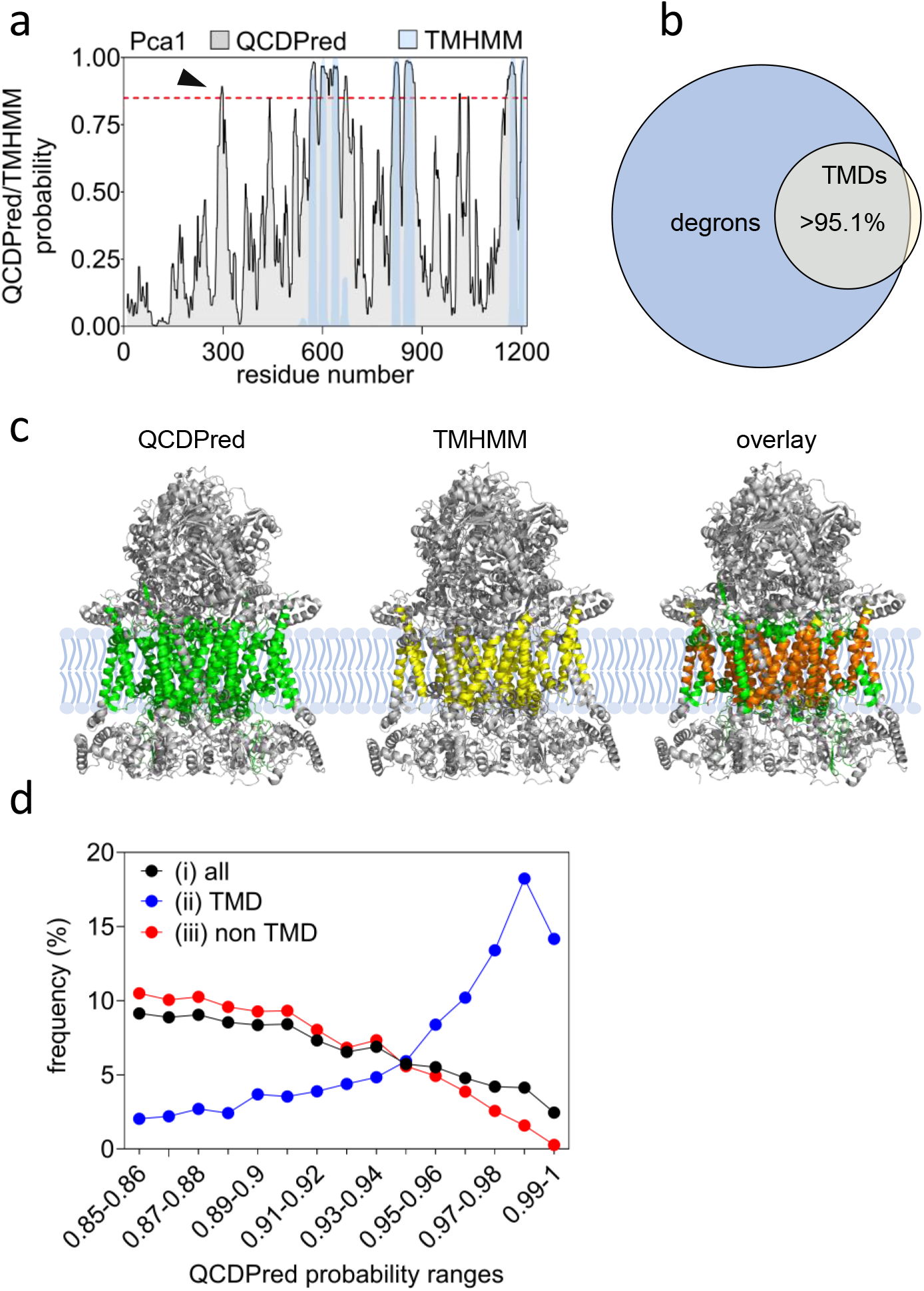
QCDPred assignment of TMDs as QCAP degrons. (a) Map of degrons prediction by QCDPred (in grey) and TMDs prediction by TMHMM (in light blue) within Pca1 sequence. Marked by an arrowhead is the Pca1 cytosolic degron. A degron cutoff probability is marked at P = 0.85 by a dashed red line. (b) Venn diagram of the relationship between yeast TMDs, extracted from the TM Helix Hidden Markov Model (TMHMM) algorithm, and QCDPred degron probabilities (P ≥ 0.85). (c) Assignment of QCDPred calculated degrons (in green) and TMHMM predicted TMDs (in yellow) to the cytochrome b-c1 complex (PDB #6T0B). overlay regions are colored in orange. (d) Frequency distribution of (i) all QCDPred assigned degrons (P ≥ 0.85), (ii) degrons composed of TMDs, and (iii) degrons that are not part of a TMD, sorted into probability ranges between 0.85 -1.

Since the QCDPred algorithm is based on data from short peptides and thereby devoid of the cellular context, we wished to empirically assess the putative function of TMDs as potential degrons in a more physiological setting. To this end, three single-pass type-I proteins were selected to examine our hypothesis. These include the COPII-coated vesicles protein Erp2 ^39^, Atg27, which is involved in autophagy and coated-vesicle transport ^40^, and Ksh1, which functions in the early steps of the secretory pathway ^41^. When expressed as GFP fusions, all three proteins exhibited membrane localization (Figure 4a). QCDPred assigned degron function to both the N-terminal signal peptides (SPs) that mediate ER insertion ^42^ and the TMDs of the three proteins (Figure 4b). This was confirmed experimentally by fusing the TM or SP regions of the three proteins C-terminally to GFP (a position that is unlikely to support ER translocation), followed by flow cytometry analysis (Figures 4c, 4d). Both TMs and SPs are proteasome substrates, as demonstrated in cells expressing the corresponding plasmids that were treated with BZ (Supplementary Figure 5a). Thus, besides ER targeting, SPs may additionally function as QCAP degrons if translocation fails.

**Figure 4.**
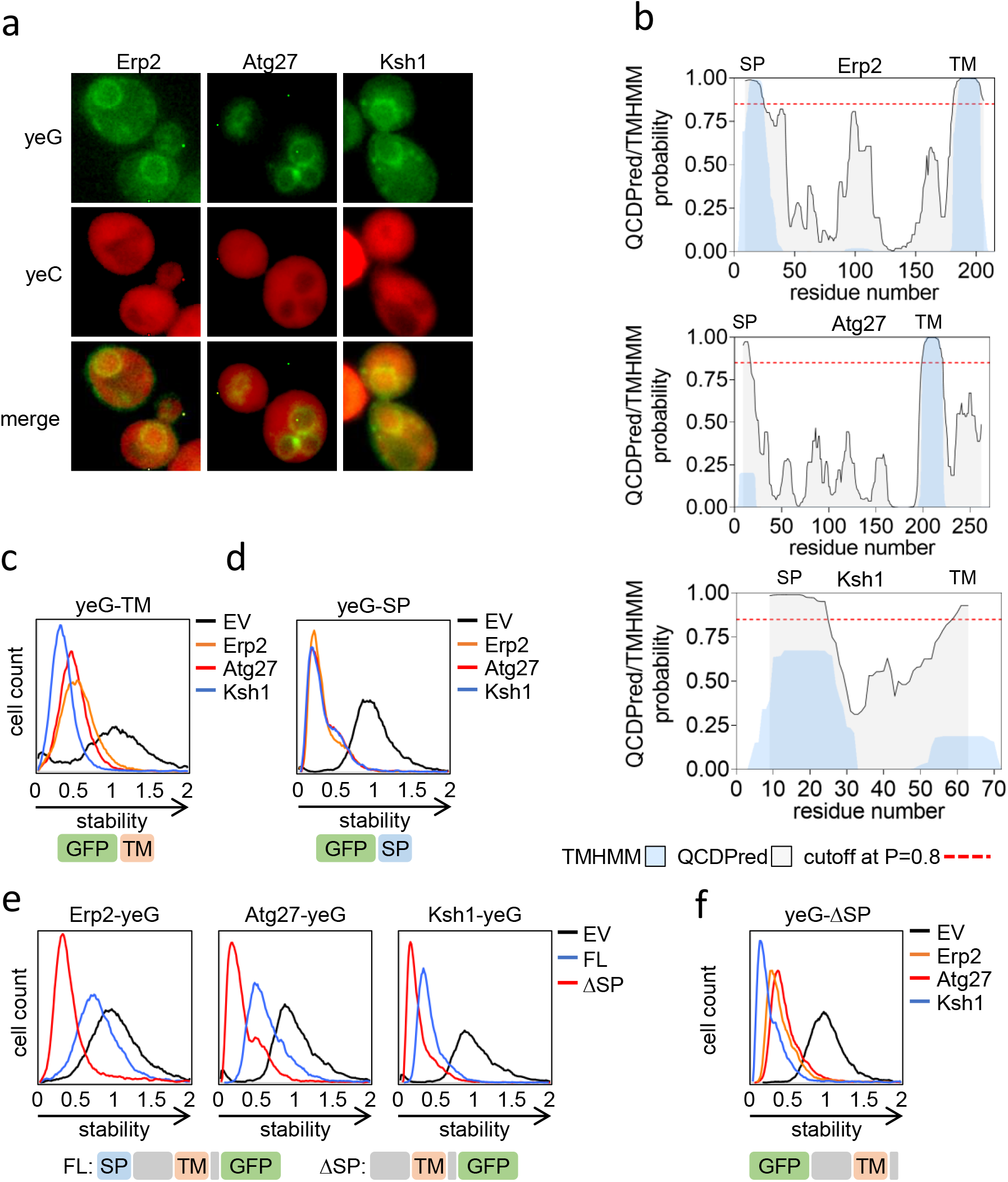
Transmembrane domains function as QCAP degrons. (a) Membrane localization of single-pass type-1 TM proteins, Erp2, Atg27, and Ksh1, determined by fluorescence microscopy. (b) Map of QCDPred and TMHMM predictions of Erp2, Atg27, and Ksh1 degron and TMD probabilities, respectively. A degron cutoff probability is marked at P = 0.85 by a dashed red line. (c, d, e, f). Flow cytometry histograms of the normalized yeG/yeC ratio in cells expressing (c, d) C-terminal appended TMDs (c) or SPs (d), (e) N-terminal appended, full length and ΔSP proteins, and (f) C-terminal appended ΔSP proteins. Illustrated below each panel are the composition and domain order of the GFP-appended proteins. TM: transmembrane, SP: signal peptide. Scale: Median value of the yeG/yeC ratio in empty vector (EV) control was set as one. All other histograms were distributed accordingly.

As the removal of the SP would likely disrupt ER targeting, we hypothesized that this would in turn lead to a rapid turnover of TMD-containing proteins via recognition of TMDs as degrons. This was tested by determining the steady-state levels, with or without the SP, of Erp2, Atg27, and Ksh1 (Figure 4e, Supplementary Figure 5b). Appending the full-length proteins N-terminally to GFP resulted in partial destabilization, assumingly by QCAP, that indeed, was greatly enhanced by the removal of the SP. Curiously, _∆SP_Atg27 did not respond to proteasome inhibition while levels of _∆SP_Erp2 and _∆SP_Ksh1 were significantly increased (Supplementary Figure 5b), suggesting a proteasome-independent degradation mechanism for the mislocalized autophagy-associated protein. Notably, positioning of the cytosolic exposed TMD degrons is immaterial to their function as both N-terminally-and C-terminally-appended SP-excluded proteins were substantially unstable (Figures 4e, 4f).

### QCAP degron characteristics

The data so far indicate that in the yeast proteome QCAP degrons are widespread and that these regions are prevalent in hydrophobic residues while negatively charged residues are depleted. Nevertheless, we found that within hydrophobic degrons there is a bias toward specific residues (Figure 5a): Comparing the distribution of amino acids in hydrophobic TMDs with high degron probability (P ≥ 0.85) to a small set of TMDs with low degron probability (P < 0.85), we found that the former is enriched in bulky and branched hydrophobic amino acids, while the latter express small, non-polar, amino acids instead. In addition, we identified a prevalence of alpha-helical configurations in QCAP degrons in their native protein context (Figure 5b). This not only agrees with the established helical structure of TMDs but also correlates strongly with an increase in the probability of non-TMD degrons (Figure 5c). Interestingly, PQC degrons were hardly found in N/A (not assigned) regions that are likely intrinsically disordered (Figure 5b). This implies that for most QCAP degrons to become active, the protein must be structurally perturbed so that the degron is exposed. Indeed, we have found that for disordered proteins and regions there is a correlation between the presence of predicted degrons and the abundance and half-lives of the proteins ^24^. Altogether, our data indicate that QCAP degrons are enriched in bulky hydrophobic TMD-like entities.

**Figure 5.**
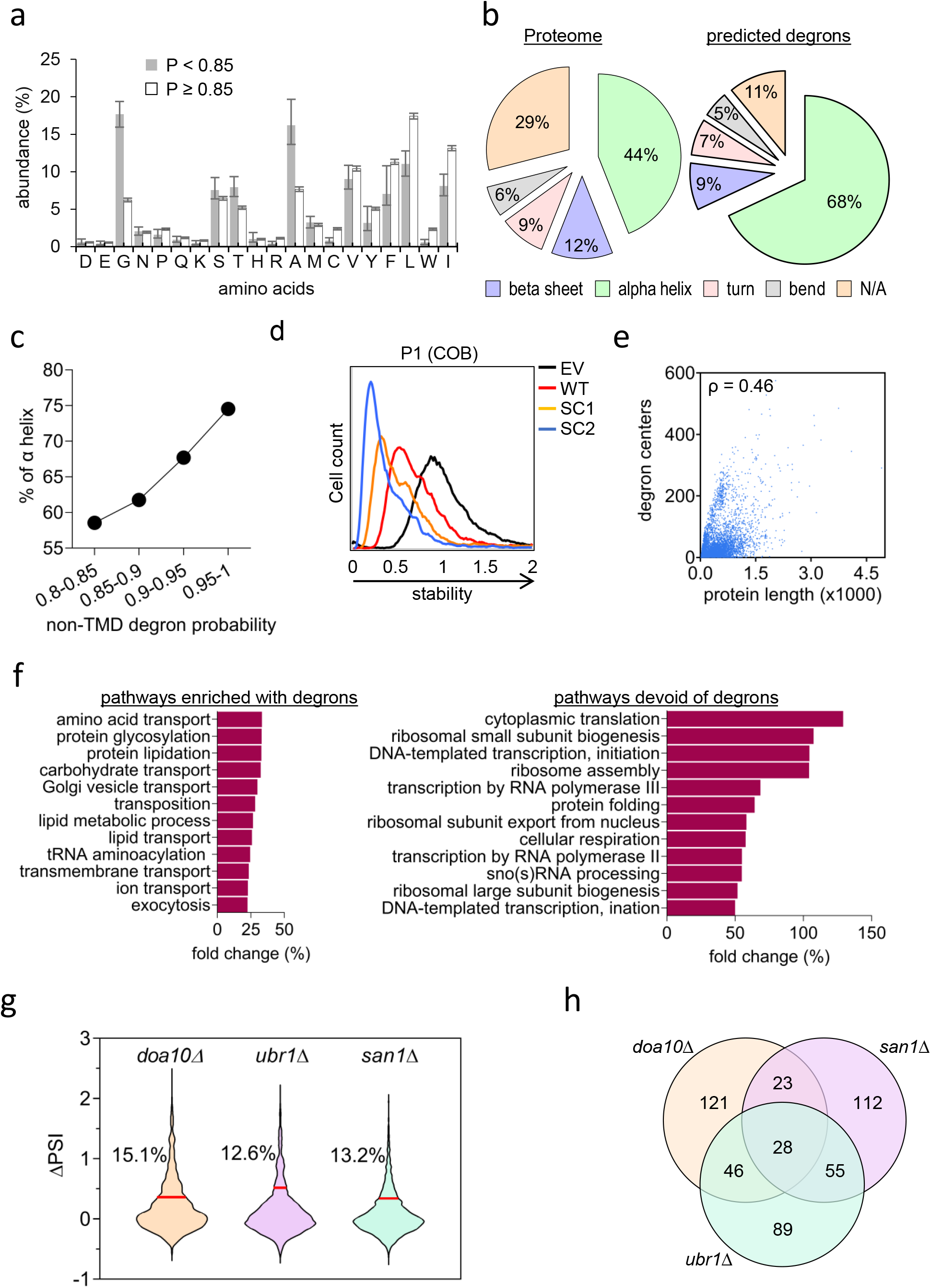
Characterization of QCAP degrons. (a) Comparison of mean amino acid distribution between TMDs with high degron probability (P ≥ 0.85, N=6,064) to those with lower probability (P < 0.85, N=107). Error bars represent low and high confidence intervals. (b) Pie chart of the classification and relative proportions of protein secondary structure within the yeast proteome versus that of QCDPred-predicted degrons, based on the AlphaFold Protein Structure Database. (c) Plot of the percentile of α helix structures versus QCDPred assigned non-TMD degron probabilities divided into four equal bins. (d) Flow cytometry histograms of the normalized yeG/yeC ratio in cells expressing P1 peptide emerged from the yeast protein COB and two randomly scrambled variants SC1 and SC2. Scale: Median value of the yeG/yeC ratio in empty vector (EV) control was set as one. All other histograms were distributed accordingly. (e) Plot of QCDPred calculated degron centers, as a function of protein length. ρ: Spearman’s correlation coefficient. P_value_ < 0.0001. (f) Gene ontology (GO) process annotations of the fold change of top twelve significant pathways (P-value < 0.05) enriched or devoid of degrons, compared to a reference yeast proteome (strain S288C). (g) Violin plot displaying changes in the PSI of 2175 degrons (PSI ≤ 1.7) upon knocking out ORFs of the tested QCAP E3 enzymes. ∆PSI values between degrons in *pdr5∆* strain (PSI < 1.7, 2175 peptides) *and E3∆* strains were calculated. The red line marks two standard errors from the mean for each strain. Degrons above this threshold were considered as stabilized by the knockout. The Percentile of stabilized degrons is indicated for each tested QCAP E3. (h) Venn diagram displaying overlapping functions of the tested QCAP E3 enzymes. Sequences of the top 10% ∆PSI values for each knockout strain (218 peptides) were compared.

We next examined the role of the linear order of amino acids within a degron on its function. To this end, the amino acid sequence of a peptide P1 from the yeast protein COB of the b-c1 complex was scrambled to form two random peptides having identical amino acid content, hence also identical degron probability when predicted by QCDPred (Table 1). When fused to the C-terminus of GFP, all three peptides were predicted to form helices, however to various degrees (Supplementary Figure 6). Both scrambled peptides not only conferred GFP degradation but were even more potent than the original P1 peptide, indicating that the general chemical properties of the degron, rather than its exact linear sequence is a principal QCAP degron determinant (Figure 5d). The results, however, also show that—at least in this case—the patterning and sequence can play a modulatory role of the degron strength.

### Degron presence correlates with proteins function

We next searched for roles governing the distribution of degrons in the proteome. Specifically, whether it is simply dependent on the probability of finding a defined sequence within the entire proteome or associated with specific protein property or function. As shown in Figure 5e, a weak monotonic relation was observed between protein length and degron presence (Spearman rank correlation coefficient (ρ) of 0.46), thus excluding random distribution of the QCAP degronome. We then analyzed degrons through Gene Ontology (GO) annotation (Figure 5f). To this end, the yeast proteome was classified, based on QCDPred score into two groups — one with and one without degrons at high significance (P ≥ or < 0.85). Each group was then used as input for determining GO processes using the Saccharomyces Genome Database (SGD) Gene Ontology Slim Term Mapper (https://www.yeastgenome.org/goSlimMapper) Data in Figure 5f and Supplementary Figure 7 show that proteins engaged in transport and lipid metabolism are enriched in QCAP degrons. Many of these protein classes are integrated into membranes, an observation that agrees with our finding that TMDs can also act as QCAP degrons when exposed to the PQC system. Conversely, proteins, where QCAP degrons are underrepresented, are mostly involved in transcription, translation, ribosome assembly, as well as protein folding. The latter group is of particular interest as it implies that the exclusion of QCAP degrons in chaperones involved in proteolysis renders them resistant to degradation themselves. Indeed, QCDPred analysis of a collection of cytosolic/nuclear yeast chaperones predicted one or more degrons in Hsp90, Hsp104, and Hsp110 family members, while Hsp40 and Hsp70 family members, that are directly involved in proteolysis ^2^, were mostly devoid of QCAP degrons (Supplementary Figure 8).

### Partial selectivity of QCAP E3 ligases

To learn about UPS-dependent QCAP functionality, we next examined degron specificity of Ubiquitin-protein E3 ligases. To this end, the aforementioned peptide library was inserted into yeast strains lacking one of three well-defined QCAP E3 ligases — Doa10, which is a multi-TMDs E3 ligase that localizes to the outer leaflet of the ER membrane and the nuclear envelope ^43^, Ubr1, which is a soluble protein residing in the cytoplasm and the nucleoplasm ^44^, and San1, which is exclusively nuclear E3 ^45^. yGPS-P_lib_ transformation into E3 deleted (*E3∆*) strains was followed by FACS and NGS, and PSI scores of degrons having P < 1.7 were determined and compared to that of *pdr5∆* cells that served as a control strain (Supplementary Data 2). All knockout groups displayed a significant increase in degrons PSI scores (Kruskal-Wallis test P < 0.001). A violin plot of the change in PSI score (ΔPSI = PSI_E3∆_-PSI_WT_) indicates that Doa10 substrates are the largest group of degrons in the tested peptidome (Figure 5g). We note, however, that because the PSI scale is effectively determined by the distribution of degron potential within an individual experiment, it is difficult to interpret ΔPSI scores on an absolute scale. A Venn diagram determining E3 functional overlaps indicates that approximately one-third of degrons were recognized by two or more E3 enzymes (Figure 5h). This finding is consistent with that of Hickey and colleagues who showed distinct yet overlapping QCAP E3 ligases substrate specificity governed by E3 subcellular localization ^18^.

### Conserved constraints governing degron potency

Considering the high conservation of QCAP pathways in the evolution of all eukaryotes, we hypothesized that QCAP degron properties are similarly well preserved. To test this paradigm, we next investigated how well the yeast-based QCDPred algorithm predicts the presence of QCAP degrons in other organisms and selected the influenza C virus p42 protein and human serum and glucocorticoid-inducible kinase 1 (SGK1) as test cases. The influenza p42 contains a signal peptidase site at residue 259 that upon cleavage yields the p31 and CM2 proteins (Figure 6a, *upper panel*). CM2 integrates into the ER membrane through a single TMD, while p31 is rapidly degraded by the UPS via a degron at the C-terminal region ^46^. As anticipated, both the C-terminal region of p31 and the TMD of p42 were predicted by QCDPred to function as degrons (Figure 6a, *lower panel*), however, during viral infection only the C-terminal region of p31 likely functions as a degron because it is accessible to the degradation system. Furthermore, Arteaga and co-workers have previously demonstrated that an amphipathic helix at the N-terminus of SGK1 targets the protein for proteolysis ^47^. Indeed, QCDPred analysis of SGK1 revealed three potential degrons, the strongest of which is placed between amino acids 17 and 29, the same region that was previously identified as a QCAP degron ^47^ (Figure 6b).

**Figure 6.**
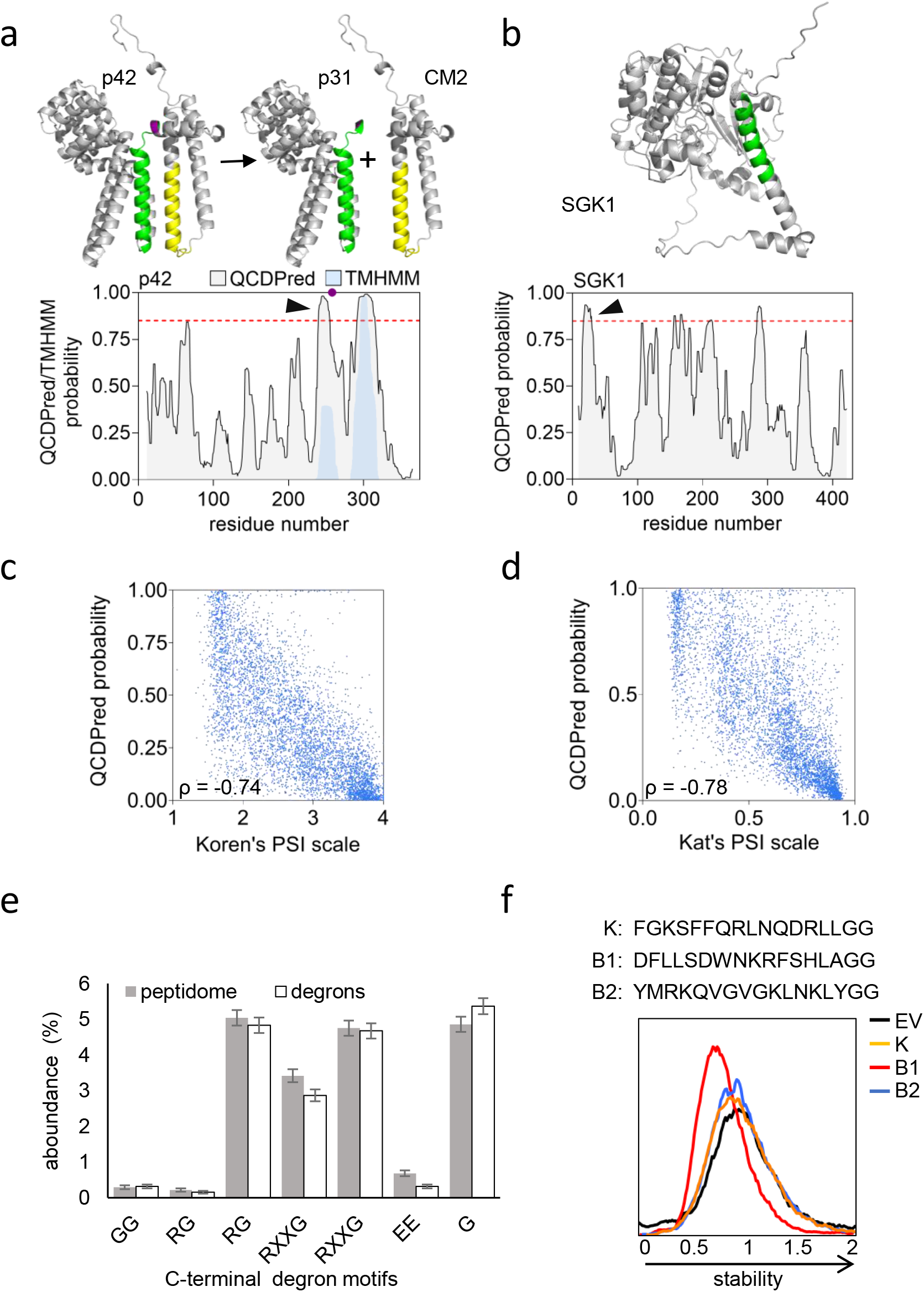
Degron projections in non-yeast organisms. (a) *upper panel* Assignment of QCDPred projected degrons into the influenza C virus Polyprotein p42 trRosetta-predicted 3D structure (Template modeling score: 0.642). Shown are the schematics of p42 cleavage at an internal SP site (marked in purple) into p31 and CM2. *Lower panel*: QCDPred and TMHMM probabilities of p42 degrons (in grey) and TMDs (in light blue), respectively. *Arrowhead* indicates the position of the postulated p31 degron. ‘A degron cutoff probability is marked at P = 0.85 by a dashed red line. (b) *Upper panel*: Assignment of a QCDPred-identified QCAP degron into the AlphFold-predicted SGK1 structure (#AF-O00141-F1). *Lower panel*: QCDPred-calculated degron probabilities within SGK1. *Arrowhead* indicates the position of the postulated SGK1 degron. A degron cutoff probability is marked at P = 0.85 by a dashed red line. (c) Scatter plot comparing PSI values of 23mer C-terminal peptides of the human proteome ^22^ and averaged QCDPred assessment of the same fragments. Prediction scores were averaged between 7 different amino acid centers within the 23 amino acids length of the original peptides. Shown are values of random 25% peptides from Koren’s screen. ρ: Spearman’s correlation coefficient. P_value_ < 0.0001. (d) Scatter plot comparing PSI values of 15mer N-terminal peptides of the yeast *Saccharomyces cerevisiae* proteome ^48^ and QCDPred assessment of the same fragments. ρ: Spearman’s correlation coefficient. P_value_ < 0.0001. (e) Comparison of the mean distribution of the six principal C-terminal degradation motifs identified by Koren at al. ^22^, between the tested peptidome and calculated degrons (PSI ≤ 1.7). Error bars represent the standard deviation. (f) Flow cytometry histograms of the normalized yeG/yeC ratio in cells expressing fused peptides with a GG end: K-a C-terminal degron identified by Koren at al. ^22^, and B1-2 from the current screen. Scale: Median value of the yeG/yeC ratio in empty vector (EV) control was set as one.

Having demonstrated that the tested non-yeast QCAP degrons can be predicted by QCDPred, we next wanted to investigate the universality of QCAP degrons by testing whether the principles established for yeast degrons generally apply to the human degronome. To this end, we assessed the correlation between PSI values, previously assigned by Koren et al., for C-terminal regions of the entire human proteome ^22^, and their average QCDPred probabilities. The comparison yielded a high correlation (ρ = -0.74) (Figure 6c), similar to that observed by Kats et al., for the yeast N-terminome (ρ = -0.78) ^48^ (Figure 6d), indicating that indeed principles of QCAP degron features are evolutionarily conserved. Consequently, average QCDPred scores were assigned to each amino acid in both the yeast and human proteomes (Supplementary Data 3,4) and we also provided QCDPred as a web-server tool to predict degrons ^24^.

Koren et al. have shown that C-terminal glycine residues are underrepresented in the eukaryotic proteome and proposed that the depletion of glycine at the C termini of eukaryotic proteins is a result of avoidance of E3s targeting glycine-end degrons ^22^. Glycine residues are equally underrepresented in *S. cerevisiae*, which implies for a similar role in degradation (Supplementary Figure 9). To examine the possible role of glycines and other C-terminal degradation motifs described by Koren et al., in *S. cerevisiae*, we analyzed their abundance at the C-terminus of high confident degrons within the tested degronome (PSI ≤ 1.62) and compare it to that of the entire peptidome. Surprisingly, no significant difference between the two groups was observed (Figure 6e), suggesting that, unlike in humans, C-terminal degrons do not play a substantial role in determining the half-lives of *S. cerevisiae* proteins. In agreement with these findings, neither a degron peptide from the Koren screen (K) nor re-cloned peptides from the peptidome, having two glycine residues at their C-termini, conferred GFP destabilization in yGPS-P vector (Figure 6f).

## Discussion

A major barrier to unearthing QCAP degrons has been their unconformity. Therefore, authentic degron discovery largely relied on the screening of peptide libraries in search of sequences that induce the degradation of otherwise stable proteins and then, trying to deduce consensus sequence requirements. ^15^ Indeed, these efforts led to the discovery of many artificial and physiological degrons but were still far from distinguishing common degradation motifs in QCAP. A breakthrough in degron discovery has been provided by the development of the pioneering GPS-peptidome methods to study protein degradation and identify degron motifs ^22,23,49^. By implementing GPS-P in yeast we have here identified a large cohort of authentic degrons that was subsequently used to train a degron prediction algorithm ^24^. Thus, this large group of peptides that determine a variety of half-lives, provide consensus degron features that are compatible with TMD properties. Further functional validation established TM-like degron features as key determinants of QCAP and possibly of the proteome half-life.

The discovery that E3 ligases selectively bind substrates through recognition of distinct determinants (degrons) established their role as substrate recognition modules of the UPS ^50^. Consequently, the identification of degrons has become one of the focal points of UPS research. Initially, the hunt for degrons identified mostly acquired determinants that are, for example, the result of transient post-translational modifications such as phosphorylation that induce timely, regulated degradation via dedicated E3 ligases ^51^. Obviously, acquired degrons do not account for the majority of QCAP, carried out by specialized E3 ligase systems that monitor protein folding state, presumably recognizing internal sequences that may become exposed following conformational aberrations. This assertion was confirmed during the characterization of *Deg1* degron of the Doa10 E3 ligase ^52^. *Deg1* is masked in the Mata1/Mat α2 mating-type dimer and exposed upon complex dissociation ^53^. Critical for *Deg1* degron function is an amphipathic helix ^53^, also present in other synthetic and authentic Doa10 substrates ^27,28,47,54^. Despite hydrophobicity constituting a primary Doa10-recognition determinant ^18^, not all QCAP degrons conform to the same consensus ^19^ (see a comparison in Supplementary Data 5). Overall, the initial degron discoveries, while gaining important insights into degron complexities, could not account for the entire QCAP.

At present, hydrophobic stretches are considered key determinants of QCAP in all eukaryotes, from yeast to mammals ^18^. Our observations that QCAP degrons are to a large extent determined by amino acid composition implies a broad degron recognition mechanism. The prevalence of QCAP degrons function was previously demonstrated by overlapping degron recognition by E3 ligases (Figure 5g and ^17^). As protein structure is dynamic, QCAP sensing of protein folding state via the exposure of hydrophobic, TM-like regions and their nonselective E3 ligase recognition can now provide a plausible mechanism for regulation of the proteome stability. As E3 ligases are rate-limiting for ubiquitin-dependent degradation, recognition of exposed hydrophobic stretches— directly or indirectly—by multiple E3 ligases may increase cellular degradation capacity in response to diverse stress conditions where aberrant protein overload might lead to proteotoxicity^55^.

That TMDs can operate as degrons is not surprising as membrane-embedded sequences within integral ER-membrane proteins have already been shown to display a degron function. These include the C-terminal TMD of the T-cell receptor α subunit (TCR α) ^56,57^ as well as other lone proteins that are normally part of TM protein complexes ^58,59^. Single and homomeric ER-embedded proteins, such as the E2 enzyme Ubc6 and the C-terminal TMDs of yeast and human HMG CoA reductase can similarly undergo QCAP via their TMDs that act as degrons ^60–62^. By demonstrating that single-pass TM proteins devoid of their SPs were rapidly degraded by the UPS (Figures 4e, 4f), we have expanded this view by establishing TMDs as conserved QCAP degrons of non-integrated TM proteins.

Our findings that TMDs can act as degrons are compatible with a pre-insertion degradation mechanism, operating at the cytosolic side of the ER membrane, that subjects Signal Recognition Particle (SRP)-independent substrates of the glycosylphosphatidylinositol (GPI) anchored proteins to QCAP ^63^. It is also compatible with SRP-independent insertion mechanisms of Atg27 and Ksh1 ^64^ that eliminate subpopulations that may evade the secretory pathway (Figure 4f). Moreover, in mammals, BAG6 and its associated protein UBQLN4 were shown to recognize the exposed hydrophobicity of TMDs of proteins that evade the secretory pathway and trigger their proteasomal degradation ^65,66^. Thus, exposure of both the SP and TMDs may ensure that suboptimal or complete failure of ER membrane integration would result in QCAP. Failure to degrade mislocalized membrane proteins may result in cytotoxicity, due to enhanced formation of insoluble intracellular bodies or aberrant insertion in the mitochondria membrane ^64^. Our observation that SP-devoid integral membrane proteins are subjected to rapid degradation (Figures 4e, 4f) is fully compatible with this assumption.

Molecular chaperones that discern misfolded proteins also participate in misfolded protein degradation ^2,67^. Auxiliary PQC chaperone functions include substrate solubilization, mediating E3 binding, as well as delivery of ubiquitinated proteins to the proteasome. The prevailing view of chaperone function in QCAP asserts that Hsp70s initially recognize misfolded substrates and deliver them to an E3 ligase, a function facilitated by Hsp40s and nuclear exchange factors that catalyze ATP hydrolysis and ADP exchange, respectively ^68^. While a role for Hsp70s/Hsp40s in misfolded substrates targeting the human E3 ligase, the carboxy-terminus of Hsc70 interacting protein (CHIP), is well established ^69^, whether Hsp70s/Hsp40s similarly mediate the recognition of QCAP degrons by their cognate E3s remains to be determined.

Surprisingly, despite our observation that glycine residues are underrepresented at the C-termini of the *S. cerevisiae* proteome, sequence-specific C-terminal degrons do not seem to play a principal role in yeast protein turnover determination. One explanation for the discrepancy between the human and yeast proteomes is that yeast does not encompass Cullin Ring Ligases (CRLs) that recognize C-terminal degrons. Indeed, the principal CRLs that take part in C-terminal degron recognition, namely Cul2 and Cul4 family members are absent in *S. cerevisiae* and the relevant F-boxes are also missing. Alternatively, C-terminal yeast degrons may be recognized by other E3 ligases with more complex specificity. Perhaps the role of C-terminal glycines in fungal protein degradation is more context-dependent, e.g., that it requires other, more distant degron elements that are not present in the 17mer peptides. further studies of the stability of yeast proteome are required in order to identify and characterize these speculative distal elements.

## Conclusion

Adaptation of the GPS-P technology into yeast research facilitated the identification of QCAP degron sequences within the eukaryotic peptidome. The substantial number of identified degron sequences thus enabled a machine-learning algorithm to faithfully define consensus degron features. The ability of QCDPred to identify degrons in viral and human proteins (Figure 6), supports the notion that the majority of the proteome contains QCAP degrons with conserved characteristics.

The extent of TM-like degron function remains to be determined. Whether it is the “Holy Grail” that controls proteome stability or if other degron modules also play a substantial role remains to be elucidated. However, the properties of TM-like domains and their conditional exposure can now, for the first time, provide an expansive explanation of QCAP function in the maintenance of the eukaryotic proteome.

## Acknowledgement

We are grateful to Prof. Richard Kulka for his inspiration and support. We also thank Dr. Yuval Reiss for diligently reviewing the manuscript and Dr. Itay Koren for advice during the setup of yGPS-P. B.M. acknowledges the support of the Neubauer doctoral fellowship fund. This work was supported by NSF-BSF grant 2016722 (to R.G.G and T.R) and the Novo Nordisk Foundation centre PRISM (NNF18OC0033950; to R.H.-P. and K.L.-L.).

## Author contributions

Conceptualization, B.M., R.G.G. and T.R.; Methodology, B.M., R.G.G., T.R., K.E.J, K.L.L. R.H.P; Software, K.E.J., K.L.L., S.A., A.C., and N.F.; Validation, B.M.; Formal Analysis, K.E.J., and S.A.; Investigation, B.M., S.A., K.E.J., and T.R.; Resources, B.M., S.A., K.L.L.; Writing-Original Draft, B.M., S.A., and T.R.; and Writing-Review & Editing, K.E.J, K.L.L., R.H.P., R.G.G., N.F., and T.R.

## Materials and Methods

### Parental plasmid for yGPS-P screen

Plasmid pGADT7-ADH700-yeCherry-p150-yeGFP-DHFR was obtained from Addgene (#24378) ^25^ and was used as a template for PCR cloning of ADH700-yeCherry-P150-yeGFP into pTR1412 ^16^ at NotI and XmaI restriction sites. A 5-mer linker was added downstream to GFP to create the parental plasmid pTR1861 for yGPS-P screen (Figure 1a). pTR2089, the parental plasmid for yGPS-P N-terminal cloning, was constructed by overlap extension PCR ^70^, producing a fragment containing PacI and BamHI restriction sites, that was placed upstream to GFP in pTR1861 by ligation.

### Cloning and mutagenesis

Plasmids used in this study are listed below. DNA fragments encoding the full-length, ∆SP, TMD only, and SP only versions of Erp2, Atg27, and Ksh1 were amplified from genomic DNA of *wild-type* yeast strain BY4741. The PCR fragments were subjected to digestion with restriction enzymes XmaI and NheI (for C-terminal cloning) or with PacI and BamHI (for N-terminal cloning). The resulting fragments were subcloned into pTR1861 or pTR2089, respectively.

Oligos insertions into pTR1861 were done by heating a single DNA pair containing the wanted insertion (see “Oligos Table” for details) and flanking sequences responsible for overhangs compatible with XmaI and NheI restriction sites and cooling down slowly to create double-stranded fragments. The double-strand DNA was ligated directly into pTR1861 and digested by the same restriction enzymes. Mutagenesis of peptide P3 and Pca1 was conducted using QuikChange Lightning Site-Directed Mutagenesis Kit, according to the manufacturer’s instructions (Agilent). All products were verified by sequencing.

### Generation of a peptide library (yGPS-P_lib_)

Three hundred thirty-five proteins, corresponding to 23 yeast complexes were first encoded as DNA bases using the Saccharomyces Genome Database (SGD) website. Then, DNA corresponding to the open reading frame of each protein was divided into 51 bp (17-mer) fragments with 36 bp (12-mer) overlaps between neighboring oligonucleotides (tiling). The fragments also contained two flanking 12 bp primers that match the vector sequence to enable Gibson assembly. The corresponding oligonucleotides were synthesized by LC Sciences (Houston, TX), amplified by PCR, and cloned by Gibson assembly master mix kit (New England Biolabs) into pTR1861 at XmaI and NheI restriction sites, followed by transformation into electro-competent DH10B bacterial cells. Approximately two million colonies were scraped from plates and pooled and plasmid DNA was purified using PureLink^®^ HiPure plasmid filter Midiprep kit (Invitrogen). The resulting plasmid library was transformed into TRy1392 *pdr5∆* yeast strain, followed by selection on leucine-deficient media (SD-Leu). Surviving cells were scraped, pooled, and frozen immediately in 25% glycerol at -80°C.

### Generation of a degron library (yGPS-P_deg)_

Yeast cells expressing yGPS-P_lib_ were grown O/N on SD-Leu media to mid-log phase. Cells were subjected to BD FACS Aria III instrument using 488 nm and 561 nm lasers for capturing the fluorescence emission of GFP and Cherry, respectively. One million cells having the lowest 10% yeG/yeC ratio, representing cells harboring unstable GFP, were separated (Supplementary Figure 1). Sorted cells were incubated O/N in SD-Leu media, then divided into aliquots and frozen in 25% glycerol at -80°C.

### Degron screen

Cells expressing yGPS-P_lib_ were grown to mid-log-phase and sorted by FACS BD-Facsaria III into four equal gates, each containing 2.5 million yeast cells, based on their yeG/yeC ratio (Figure 2a). Plasmids from each gate were purified using Zymoprep yeast Plasmid Miniprep II (Zymo Research).

### Preparation of plasmid DNA for NGS

DNA sequencing by NGS consisted of two PCR amplification steps with KAPA HiFi HotStart ReadyMix PCR Kit (Roche). The first step (18 cycles) was performed using primers flanking the peptides, with overhangs complementary to Illumina adapters (primers NGS-F, NGS-R). The second step was performed using standard N-series Illumina barcoded adapters (12 cycles). Sequences were size-selected using SPRI beads for NGS. Samples were subsequently pooled, purified on an agarose gel, and sequenced on an Illumina NextSeq 500 machine.

### Data pre-processing

Sequencing data were processed using a custom pipeline written for the R project for statistical computing (https://www.R-project.org). Reads were aligned to the expected oligo database with bowtie2 ^71^. Sequences corresponding to the tiled peptides were counted and assigned to the different strains based on forward and reverse barcodes.

Protein stability indices (PSIs) were calculated according to Yen et al ^21^. Briefly, the frequency, *f*_*ig*_, of peptide, *i*, in gate, *g*, was multiplied by gate number (1-4) and summed up, yielding a stability score between 1 (maximally unstable) and 4 (maximally stable)

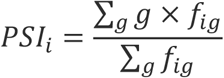

### Flow cytometry

Yeast cells were grown to mid-log-phase on SD-Leu media and analyzed on a CellStream analyzer instrument (Merck) using 488 nm and 561 nm lasers for capturing the fluorescence emission of GFP and Cherry, respectively. For each condition, 10,000 events were analyzed and presented on histogram using FlowJo software (BD Biosciences). Experiments were repeated two or more times.

### Immunoblotting

Cells were grown to mid-log-phase and pelleted by centrifugation (3,500 × g, 5 min). Cells were incubated with 0.1N NaOH for 5 min, then pelleted by centrifugation (17,000 × g, 3 min), and SDS-PAGE sample buffer containing 50 mg/ml Dithiothreitol was added, followed by boiling for 5 min. Proteins were separated on SDS-PAGE, transferred to a PVDF membrane, blocked in 10% Dry Milk in TBS + 0.1% Tween-20 (TBS-T), and then probed with primary antibodies for one hour at room temperature. Following three washes with TBS-T, the membrane was incubated with a secondary antibody for 0.5 h at room temperature, then washed three times with TBS-T. Membranes were incubated with ECL mix (Thermo Fisher Scientific) for 2 min and reactive bands were visualized using Fusion Pulse (Vilber Lourmat).

### Fluorescence Microscopy

Imaging was performed with Olympus IX71 inverted microscope with an x 60 oil objective lens. Fluorescence was excited with 576 nm for Cherry and with 488 nm for GFP. Imaging data were handled using ImageJ.

### Proteome Databases

All proteome databases were downloaded from the UniProt database server (https://www.uniprot.org) as FASTA files. These files contain the full-length protein sequence.

### PDB/CIF data

AlphaFold-2-based PDB and CIF files for single proteins were extracted from the European Bioinformatics Institute website (https://alphafold.ebi.ac.uk/). These files were used for creating a 3D protein database. Most PDB and CIF files contain information on secondary structure patterns. Each amino acid in these models was assigned a secondary structure indicator and proteome statistics were inferred.

### Transmembrane protein Data

Transmembrane protein data was extracted from the TM Helix Hidden Markov Model (TMHMM) algorithm ^72,73^, implemented with python package tmhmm.py. To compute the intersection between TMDs and degrons, each amino acid within the entire yeast proteome was evaluated for QCDPred value and TMD classification. Amino acids with QCDPred probability ≥ 0.85 that were classified as TMDs according to TMHMM were added to the intersection group.

### Antibodies used in this study

|  |  |  |
| --- | --- | --- |
| Polyclonal Rabbit anti-GFP | Abcam | ab290 |
| Monoclonal Rabbit anti-mCherry | Abcam | ab213511 |
| Monoclonal Rat anti-HA | Roche | 11867423001 |
| Monoclonal Mouse anti-PGK1 | Abcam | ab113687 |
| Polyclonal Rabbit anti-G6PD | Sigma Israel | a9521 |
| Polyclonal Rabbit anti-Ubiquitin | Dako | Z0458 |
| Goat anti-Rabbit | ImmunoResearch | 111035144 |
| Goat anti-Mouse | ImmunoResearch | 115035062 |
| Goat anti-Rat | ImmunoResearch | 112035062 |

### Chemicals

Chemicals:
| Chemical | Manufacturer | Catalog # |
| --- | --- | --- |
| Bortezomib (BZ) | A2S | A2614 |
| Cadmium | Sigma | 265365 |

### Yeast strains used in this study

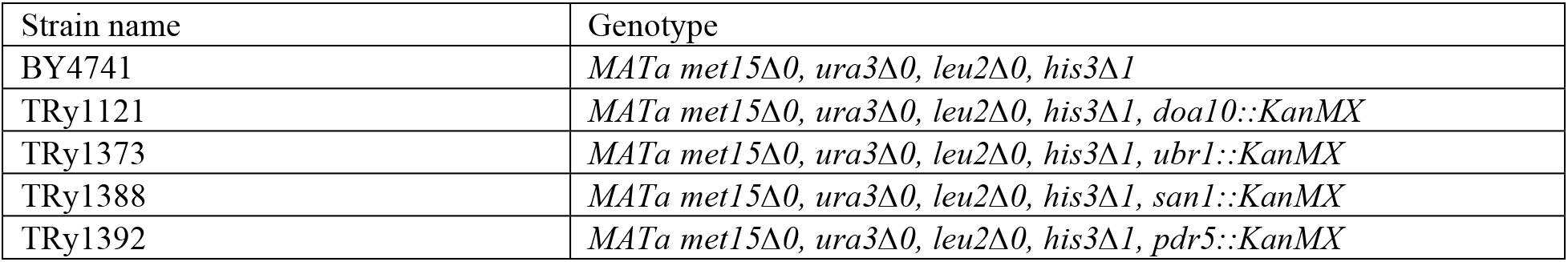

Yeast strains used in this study:
| Strain name | Genotype |
| --- | --- |
| BY4741 | *MATa met15Δ0, ura3Δ0, leu2Δ0, his3Δ1* |
| TRy1121 | *MATa met15Δ0, ura3Δ0, leu2Δ0, his3Δ1, doa10::KanMX* |
| TRy1373 | *MATa met15Δ0, ura3Δ0, leu2Δ0, his3Δ1, ubr1::KanMX* |
| TRy1388 | *MATa met15Δ0, ura3Δ0, leu2Δ0, his3Δ1, san1::KanMX* |
| TRy1392 | *MATa met15Δ0, ura3Δ0, leu2Δ0, his3Δ1, pdr5::KanMX* |

### Oligos Table

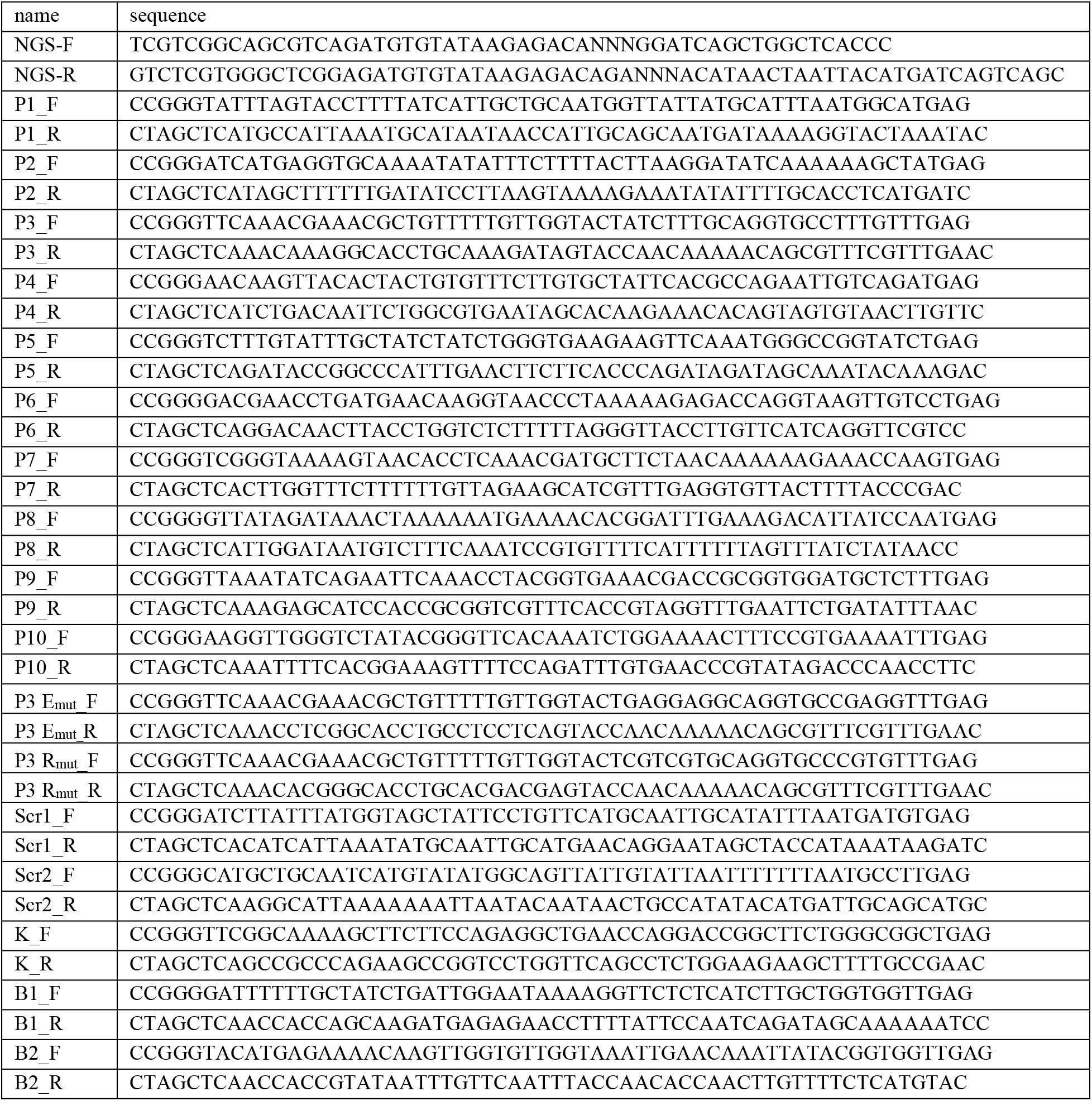

### List of plasmids used in this study

Unless indicated otherwise, clones are C-terminal fusions to yeGFP

List of plasmids used in this study Unless indicated otherwise, clones are C-terminal fusions to yeGFP
|  | Description | Additional information |
| --- | --- | --- |
| pTR1494 | pRS416-GPDp-HA-Pca1 | From Adle et al. PNAS 2009 |
| pTR1859 | pRS315-ADH700p-yeCherry-p150-yeGFP-XmaI,NheI-CYCt |  |
| pTR1861 | pRS315-ADH700p-yeCherry-p150-yeGFP-linker-XmaI,NheI-CYCt | C-terminal fusion to yeG |
| pTR1882 | pRS315-ADH700p-yeCherry-p150-yeGFP-linker-XmaI-*CL1*-NheI-CYCt |  |
| pTR1884 | pRS315-ADH700p-yeCherry-p150-yeGFP-linker-XmaI-*DegAB*-NheI-CYCt |  |
| pTR2036 | pRS315-ADH700p-yeCherry-p150-yeGFP-linker-XmaI-P1-NheI-CYCt |  |
| pTR2037 | pRS315-ADH700p-yeCherry-p150-yeGFP-linker-XmaI-P2-NheI-CYCt |  |
| pTR2038 | pRS315-ADH700p-yeCherry-p150-yeGFP-linker-XmaI-P3-NheI-CYCt |  |
| pTR2058 | pRS315-ADH700p-yeCherry-p150-yeGFP-linker-XmaI-P3<sub>EE</sub>-NheI-CYCt | TIFAGAFV → TEEAGAEV |
| pTR2060 | pRS315-ADH700p-yeCherry-p150-yeGFP-linker-XmaI-P3<sub>RR</sub>-NheI-CYCt | TIFAGAFV → TRRAGARV |
| pTR2039 | pRS315-ADH700p-yeCherry-p150-yeGFP-linker-XmaI-P4-NheI-CYCt |  |
| pTR2040 | pRS315-ADH700p-yeCherry-p150-yeGFP-linker-XmaI-P5-NheI-CYCt |  |
| pTR2041 | pRS315-ADH700p-yeCherry-p150-yeGFP-linker-XmaI-P6-NheI-CYCt |  |
| pTR2042 | pRS315-ADH700p-yeCherry-p150-yeGFP-linker-XmaI-P7-NheI-CYCt |  |
| pTR2043 | pRS315-ADH700p-yeCherry-p150-yeGFP-linker-XmaI-P8-NheI-CYCt |  |
| pTR2044 | pRS315-ADH700p-yeCherry-p150-yeGFP-linker-XmaI-P9-NheI-CYCt |  |
| pTR2045 | pRS315-ADH700p-yeCherry-p150-yeGFP-linker-XmaI-P10-NheI-CYCt |  |
| pTR2089 | pRS315-ADH700p-yeCherry-p150-PacI,BamHI-yeGFP-linker-XmaI,NheI-CYCt | N or C-terminal fusion to yeG |
| pTR2124 | pRS416-GPDp-HA-Pca1<sub>DD</sub> | LGCICNY → LDCDCDY |
| pTR2127 | pRS315-ADH700p-yeCherry-p150-yeGFP-linker-XmaI-ΔspErp2-NheI-CYCt | |
| pTR2128 | pRS315-ADH700p-yeCherry-p150-yeGFP-linker-XmaI-ΔspAtg27-NheI-CYCt | |
| pTR2129 | pRS315-ADH700p-yeCherry-p150-yeGFP-linker-XmaI-ΔspKsh1-NheI-CYCt | |
| pTR2149 | pRS315-ADH700p-yeCherry-p150-PacI-Erp2-BamHI-yeGFP-linker-CYCt | N-terminal fusion |
| pTR2151 | pRS315-ADH700p-yeCherry-p150-PacI-Atg27-BamHI-yeGFP-linker-CYCt | N-terminal fusion |
| pTR2152 | pRS315-ADH700p-yeCherry-p150-PacI-Ksh1-BamHI-yeGFP-linker-CYCt | N-terminal fusion |
| pTR2153 | pRS315-ADH700p-yeCherry-p150-PacI-ΔspErp2-BamHI-yeGFP-linker-CYCt | N-terminal fusion |
| pTR2155 | pRS315-ADH700p-yeCherry-p150-PacI-ΔspAtg27-BamHI-yeGFP-linker-CYCt | N-terminal fusion |
| pTR2156 | pRS315-ADH700p-yeCherry-p150-PacI-ΔspKsh1-BamHI-yeGFP-linker-CYCt | N-terminal fusion |
| pTR2160 | pRS315-ADH700p-yeCherry-p150-yeGFP-linker-XmaI-SP(Ksh1)-NheI-CYCt | SP of Ksh1 (aa 1-26) |
| pTR2162 | pRS315-ADH700p-yeCherry-p150-yeGFP-linker-XmaI-SP(Erp2)-NheI-CYCt | SP of Erp2 (aa 1-26) |
| pTR2164 | pRS315-ADH700p-yeCherry-p150-yeGFP-linker-XmaI-SP(Atg27)-NheI-CYCt | SP of Atg27 (aa 1-19) |
| pTR2167 | pRS416 GPD HA-Pca1<sub>RR</sub> | LGCICNY → LRCRCRY |
| pTR2171 | pRS315-ADH700p-yeCherry-p150-yeGFP-linker-XmaI-SC1-NheI-CYCt | Scrambled P1, variant #1 |
| pTR2172 | pRS315-ADH700p-yeCherry-p150-yeGFP-linker-XmaI-SC2-NheI-CYCt | Scrambled P1, variant #2 |
| pTR2213 | pRS315-ADH700p-yeCherry-p150-yeGFP-linker-XmaI-TMD(Atg27)-NheI- | TMD of Atg27 (aa 200-220) |
| pTR2214 | pRS315-ADH700p-yeCherry-p150-yeGFP-linker-XmaI-TMD(Erp2)-NheI-CYCt | TMD of Erp2 (aa 183-203) |
| pTR2215 | pRS315-ADH700p-yeCherry-p150-yeGFP-linker-XmaI-TMD(Ksh1)-NheI-CYCt | TMD of Ksh1 (aa 54-72) |
| pTR2279 | pRS315-ADH700p-yeCherry-p150-yeGFP-linker-XmaI-K-NheI-CYCt |  |
| pTR2280 | pRS315-ADH700p-yeCherry-p150-yeGFP-linker-XmaI-B1-NheI-CYCt |  |
| pTR2281 | pRS315-ADH700p-yeCherry-p150-yeGFP-linker-XmaI-B2-NheI-CYCt |  |

## Supplementary Information titles and legends

### Supplementary Figures

Supplementary Figure 1, related to Figure 1. (a) Amino acid composition of the tested peptidome compared to that of the proteome. (b) Pie chart of the classification and relative proportions of protein secondary structure within the yeast proteome versus that of the peptidome, based on the AlphaFold Protein Structure Database. (c) Isolation of top 10% degrons. Yeast cells expressing yGPS-P_lib_ Were grown to mid-log-phase, followed by FACS of top 10% degron population. One million cells comprising the degron library (yGPS-P_deg_) were collected, then grown O/N and frozen in liquid nitrogen or taken for further analysis. Illustrated is gating of the top 10% degron population out of a total of 10,000 cells.

Supplementary Figure 2. related to Figure 2. Maps of PSI values emerged from experimental data, average PSI values for each amino acid, and calculated QCDPred probabilities were plotted for each of the proteins in the screen that reach sufficient coverage. The PSI of each 17 amino acid tile was assigned to the central amino acid (black dots). An average PSI was calculated per amino acid from all tiles that cover a given position and this profile was smoothed by a five-window running median (black line). QCDPred scores were calculated for all possible tiles of the proteins and assigned to the central amino acid (except for the first and last 8 positions which are assigned 0.5) and the profile is smoothed using a five-window running median. Profiles are made for 306 proteins that have PSI values for 5 tiles or more.

Supplementary Figure 3, related to figure 2. (a) Determination of the steady-state levels of yeGFP-peptide fusions from the test set. Two peptides from each QCDPred probability group were selected for immunoblot analysis. Cells expressing the corresponding peptides were grown to log phase and lysed, followed by separation on 10% SDS-PAGE and transfer into PVDF membrane. Levels of the GFP-fused peptides were determined by immunoblotting with anti GFP Abs. (b) Group-I and group-III peptides are substrates of the proteasome. Cells expressing the related fusion peptides were incubated with DMSO (vehicle) or with 10µM BZ for four hours. The increase in normalized yeG/yeC in group-I and group-III peptides indicates proteasome-dependent degradation. No significant effect of BZ on Group-II peptides was observed. Scale: The median value of the yeG/yeC ratio in empty vector (EV) control was set as one. All other histograms were distributed accordingly.

Supplementary Figure 4, related to figure 3. Degrons in the b-c1 complex. (a) PSI values emerged from experimental data (black dots), average PSI values for each amino acid (colored in black), QCDPred probabilities (colored in red), and TMHMM prediction (colored in light blue) were plotted for each subunit of the b-c1 complex. A degron cutoff probability is marked at P = 0.85 by a dashed grey line. (b) Assignment of QCDPred projected Degrons (green) and TMHMM predicted TMDs (yellow) to TM proteins in the b-c1 3D structure (PDB #6t0b). Overlying regions are colored in orange.

Supplementary Figure 5, related to Figure 4. (a) SP and TM fusion peptides are substrates of the proteasome. Cells expressing the related fusion peptides were incubated with DMSO (vehicle) or with 10µM BZ for four hours. Scale: The median value of the yeG/yeC ratio in empty vector (EV) control was set as one. All other histograms were distributed accordingly. The increase in normalized yeG/yeC indicates proteasome-dependent degradation. (b) Immunoblot analysis of Erp2, Atg27, and Ksh1, with or without their SPs. Where indicated, 10 µM of Bortezomib (BZ) were added to cells for 4 hrs before cell harvesting. *FL*-full-length proteins. *∆SP-*proteins devoid of the SP. G6PD: Glucose-6-phosphate dehydrogenase.

Supplementary Figure 6, related to Figure 5. Structure prediction of GFP-fused peptides using trRosseta. Estimated Template Modeling scores: COB -0.856, SC1 -0.865, SC2 -0.828.

Supplementary Figure 7, related to Figure 5. Large-scale GO annotations of all significant processes (p-value < 0.05) that are either enriched or devoid of predicted degrons at P ≥ 0.85.

Supplementary Figure 8, related to figure 5. QCDPred analysis of key cytosolic/nuclear yeast chaperones. A degron cutoff probability is marked at P = 0.85 by a dashed red line.

Supplementary Figure 9, related to Figure 6. C-Terminal Glycine is depleted from the proteome of *Saccharomyces cerevisiae*. Amino acid proportions across the last ten positions of each proteome are shown. The data for each residue are centered to have a mean zero.

#### Supplementary Tables

Supplementary Table 1, related to Figure 2. Mutant P3 peptides and their QCDPred scores.

#### Supplementary Data

Supplementary Data 1, related to Figure 1. proteins in the screen and the complex they belong to.

Supplementary Data 2, related to Figure 2. PSI values, average PSI, and degron probabilities for peptides in the screen.

Supplementary Data 3, related to Figure 6. QCDPred analysis of the yeast proteome.

Supplementary Data 4, related to Figure 6. QCDPred analysis of the human proteome.

Supplementary Data 5. List of intact and mutated Doa10 degrons and their QCDPred probability score and GRAVY scores.

#### Data availability statement

The authors declare that the main data supporting the findings of this study, including experimental procedures and compound characterization, are available within the article and its Supplementary Information files, or from the corresponding author upon request. Plasmids encoding select cloning vectors have been deposited at Addgene for distribution. Custom code used to process and analyze peptide library data is available at: https://github.com/KULL-Centre/papers/tree/main/2022/degronome-Mashahreh-et-al/. QCDPred analyses of yeast and human proteomes are available in Supplementary Data 3,4, respectively. QCDPred analyses of other proteins of interest are available on a web server described by Johansson et al. ^24^.

## References

1. Balchin, D., Hayer-Hartl, M. & Hartl, F. U. In vivo aspects of protein folding and quality control. Science 353, aac4354 (2016).

2. Shiber, A. & Ravid, T. Chaperoning proteins for destruction: diverse roles of Hsp70 chaperones and their co-chaperones in targeting misfolded proteins to the proteasome. Biomolecules 4, 704–724 (2014).

3. Kästle, M. & Grune, T. Interactions of the proteasomal system with chaperones: protein triage and protein quality control. Prog Mol Biol Transl Sci 109, 113–160 (2012).

4. Kim, Y. E. et al. Molecular chaperone functions in protein folding and proteostasis. Annu Rev Biochem 82, 323–355 (2013).

5. Tyedmers, J., Mogk, A. & Bukau, B. Cellular strategies for controlling protein aggregation. Nat Rev Mol Cell Biol 11, 777–788 (2010).

6. Sontag, E. M., Samant, R. S. & Frydman, J. Mechanisms and Functions of Spatial Protein Quality Control. Annu Rev Biochem 86, 97–122 (2017).

7. Amm, I., Sommer, T. & Wolf, D. H. Protein quality control and elimination of protein waste: The role of the ubiquitin–proteasome system. Biochim Biophys Acta 1843, 182–196 (2014).

8. Varshavsky, A. Naming a targeting signal. Cell 64, 13–15 (1991).

9. Guharoy, M., Bhowmick, P., Sallam, M. & Tompa, P. Tripartite degrons confer diversity and specificity on regulated protein degradation in the ubiquitin-proteasome system. Nat Commun 7, 10239 (2016).

10. Rape, M. Ubiquitylation at the crossroads of development and disease. Nat Rev Mol Cell Biol 19, 59–70 (2018).

11. Koliopoulos, M. G. & Alfieri, C. Cell cycle regulation by complex nanomachines. FEBS J (2021) doi:10.1111/febs.16082.

12. Mészáros, B., Kumar, M., Gibson, T. J., Uyar, B. & Dosztányi, Z. Degrons in cancer. Sci Signal 10, (2017).

13. Christiano, R., Nagaraj, N., Fröhlich, F. & Walther, T. C. Global proteome turnover analyses of the Yeasts S. cerevisiae and S. pombe. Cell Rep 9, 1959–1965 (2014).

14. Eden, E. et al. Proteome half-life dynamics in living human cells. Science 331, 764–768 (2011).

15. Ella, H., Reiss, Y. & Ravid, T. The Hunt for Degrons of the 26S Proteasome. Biomolecules 9, (2019).

16. Geffen, Y. et al. Mapping the landscape of a eukaryotic degronome. Mol Cell 63, 1055–1065 (2016).

17. Breckel, C. A. & Hochstrasser, M. Ubiquitin Ligase Redundancy and Nuclear-Cytoplasmic Localization in Yeast Protein Quality Control. Biomolecules vol. 11 Preprint at https://doi.org/10.3390/biom11121821 (2021).

18. Hickey, C. M., Breckel, C., Zhang, M., Theune, W. C. & Hochstrasser, M. Protein quality control degron-containing substrates are differentially targeted in the cytoplasm and nucleus by ubiquitin ligases. Genetics 217, 1–19 (2021).

19. Maurer, M. J. et al. Degradation signals for ubiquitin-proteasome dependent cytosolic protein quality control (CytoQC) in yeast. G3: Genes, Genomes, Genetics g3. 116.027953 (2016).

20. Samant, R. S., Livingston, C. M., Sontag, E. M. & Frydman, J. Distinct proteostasis circuits cooperate in nuclear and cytoplasmic protein quality control. Nature 563, 407 (2018).

21. Yen, H.-C. C. S., Xu, Q., Chou, D. M., Zhao, Z. & Elledge, S. J. Global protein stability profiling in mammalian cells. Science (1979) 322, 918–923 (2008).

22. Koren, I. et al. The Eukaryotic Proteome Is Shaped by E3 Ubiquitin Ligases Targeting C-Terminal Degrons. Cell 173, 1622–1635 e14 (2018).

23. Lin, H. C. et al. C-Terminal End-Directed Protein Elimination by CRL2 Ubiquitin Ligases. Mol Cell 70, 602–613 e3 (2018).

24. Johansson, K. E., Mashahreh, B., Hartmann-Petersen, R., Ravid, T. & Lindorff-Larsen, K. Prediction of quality-control degradation signals in yeast proteins. bioRxiv 2022.04.06.487301 (2022) doi:10.1101/2022.04.06.487301.

25. Edwards, S. R. & Wandless, T. J. Dicistronic regulation of fluorescent proteins in the budding yeast Saccharomyces cerevisiae. Yeast 27, 229–236 (2010).

26. Gilon, T., Chomsky, O. & Kulka, R. G. Degradation signals for ubiquitin system proteolysis in Saccharomyces cerevisiae. Embo J 17, 2759–2766 (1998).

27. Gilon, T., Chomsky, O. & Kulka, R. G. Degradation signals recognized by the Ubc6p-Ubc7p ubiquitin-conjugating enzyme pair. Mol Cell Biol 20, 7214–9. (2000).

28. Furth, N. et al. Exposure of bipartite hydrophobic signal triggers nuclear quality control of Ndc10 at the endoplasmic reticulum/nuclear envelope. Mol Biol Cell 22, 4726–4739 (2011).

29. Alfassy, O. S. O. S., Cohen, I., Reiss, Y., Tirosh, B. & Ravid, T. Placing a disrupted degradation motif at the C terminus of proteasome substrates attenuates degradation without impairing ubiquitylation. Journal of Biological Chemistry 288, 12645–12653 (2013).

30. Swanson, R., Locher, M. & Hochstrasser, M. A conserved ubiquitin ligase of the nuclear envelope/endoplasmic reticulum that functions in both ER-associated and Matalpha2 repressor degradation. Genes Dev 15, 2660–74. (2001).

31. McCracken, A. A. & Brodsky, J. L. Assembly of ER-associated protein degradation in vitro: dependence on cytosol, calnexin, and ATP. J. Cell Biol. 132, 291–298 (1996).

32. Pla-Prats, C. & Thomä, N. H. Quality control of protein complex assembly by the ubiquitin–proteasome system. Trends Cell Biol (2022) doi:https://doi.org/10.1016/j.tcb.2022.02.005.

33. Padovani, C., Jevtić, P. & Rapé, M. Quality control of protein complex composition. Mol Cell (2022) doi:10.1016/j.molcel.2022.02.029.

34. Leppert, G. et al. Cloning by gene amplification of two loci conferring multiple drug resistance in Saccharomyces. Genetics 125, 13–20 (1990).

35. Adle, D. J. & Lee, J. Expressional control of a cadmium-transporting P1B-type ATPase by a metal sensing degradation signal. J Biol Chem 283, 31460–31468 (2008).

36. Varadi, M. et al. AlphaFold Protein Structure Database: massively expanding the structural coverage of protein-sequence space with high-accuracy models. Nucleic Acids Res 50, D439–D444 (2022).

37. Jumper, J. et al. Highly accurate protein structure prediction with AlphaFold. Nature 596, 583–589 (2021).

38. Trumpower, B. L. Cytochrome bc1 complexes of microorganisms. Microbiol Rev 54, 101–129 (1990).

39. Marzioch, M. et al. Erp1p and Erp2p, partners for Emp24p and Erv25p in a yeast p24 complex. Mol Biol Cell 10, 1923–1938 (1999).

40. Yen, W.-L., Legakis, J. E., Nair, U. & Klionsky, D. J. Atg27 is required for autophagy-dependent cycling of Atg9. Mol Biol Cell 18, 581–593 (2007).

41. Wendler, F. et al. A genome-wide RNA interference screen identifies two novel components of the metazoan secretory pathway. EMBO J 29, 304–314 (2010).

42. Hegde, R. S. & Keenan, R. J. The mechanisms of integral membrane protein biogenesis. Nat Rev Mol Cell Biol 23, 107–124 (2022).

43. Deng, M. & Hochstrasser, M. Spatially regulated ubiquitin ligation by an ER/nuclear membrane ligase. Nature 443, 827–831 (2006).

44. Prasad, R., Xu, C. & Ng, D. T. W. Hsp40/70/110 chaperones adapt nuclear protein quality control to serve cytosolic clients. J Cell Biol 217, 2019–2032 (2018).

45. Gardner, R. G., Nelson, Z. W. & Gottschling, D. E. Degradation-mediated protein quality control in the nucleus. Cell 120, 803–815 (2005).

46. Pekosz, A. & Lamb, R. A. Identification of a membrane targeting and degradation signal in the p42 protein of influenza C virus. J Virol 74, 10480–10488 (2000).

47. Arteaga, M. F. M. F., Wang, L., Ravid, T., Hochstrasser, M. & Canessa, C. M. C. M. An amphipathic helix targets serum and glucocorticoid-induced kinase 1 to the endoplasmic reticulum-associated ubiquitin-conjugation machinery. Proceedings of the National Academy of Sciences 103, 11178–11183 (2006).

48. Kats, I. et al. Mapping Degradation Signals and Pathways in a Eukaryotic N-terminome. Mol Cell 70, 488–501 e5 (2018).

49. Tokheim, C. et al. Systematic characterization of mutations altering protein degradation in human cancers. Mol Cell 81, 1292-1308.e11 (2021).

50. Hershko, A., Heller, H., Eytan, E. & Reiss, Y. The protein substrate binding site of the ubiquitin-protein ligase system. J Biol Chem 261, 11992–9. (1986).

51. Zheng, N. & Shabek, N. Ubiquitin Ligases: Structure, Function, and Regulation. Annu Rev Biochem 86, 129–157 (2017).

52. Hochstrasser, M. & Varshavsky, A. In vivo degradation of a transcriptional regulator: the yeast a2 repressor. Cell 61, 697–708 (1990).

53. Johnson, P. R., Swanson, R., Rakhilina, L. & Hochstrasser, M. Degradation signal masking by heterodimerization of MATa2 and MATa1 blocks their mutual destruction by the ubiquitin-proteasome pathway. Cell 94, 217–227 (1998).

54. Sadis, S., Atienza Jr., C. & Finley, D. Synthetic signals for ubiquitin-dependent proteolysis. Mol Cell Biol 15, 4086–4094 (1995).

55. Buchberger, A., Bukau, B. & Sommer, T. Protein Quality Control in the Cytosol and the Endoplasmic Reticulum: Brothers in Arms. Mol Cell 40, 238–252 (2010).

56. Bonifacino, J. S., Suzuki, C. K. & Klausner, R. D. A peptide sequence confers retention and rapid degradation in the endoplasmic reticulum. Science 247, 79–82 (1990).

57. Yu, H., Kaung, G., Kobayashi, S. & Kopito, R. R. Cytosolic degradation of T-cell receptor alpha chains by the proteasome. J. Biol. Chem. 272, 20800–20804 (1997).

58. Habeck, G., Ebner, F. A., Shimada-Kreft, H. & Kreft, S. G. The yeast ERAD-C ubiquitin ligase Doa10 recognizes an intramembrane degron. J Cell Biol 209, 261–273 (2015).

59. Natarajan, N., Foresti, O., Wendrich, K., Stein, A. & Carvalho, P. Quality Control of Protein Complex Assembly by a Transmembrane Recognition Factor. Mol Cell 77, 108-119.e9 (2020).

60. Walter, J., Urban, J., Volkwein, C. & Sommer, T. Sec61p-independent degradation of the tail-anchored ER membrane protein Ubc6p. Embo J 20, 3124–31. (2001).

61. Gardner, R. G. & Hampton, R. Y. A ‘distributed degron’ allows regulated entry into the ER degradation pathway. Embo J 18, 5994–6004 (1999).

62. Ravid, T., Doolman, R., Avner, R., Harats, D. & Roitelman, J. The ubiquitin-proteasome pathway mediates the regulated degradation of mammalian 3-hydroxy-3-methylglutaryl-coenzyme A reductase. Journal of Biological Chemistry 275, 35840–35847 (2000).

63. Ast, T., Aviram, N., Chuartzman, S. G. & Schuldiner, M. A cytosolic degradation pathway, prERAD, monitors pre-inserted secretory pathway proteins. J Cell Sci 127, 3017–3023 (2014).

64. Costa, E. A., Subramanian, K., Nunnari, J. & Weissman, J. S. Defining the physiological role of SRP in protein-targeting efficiency and specificity. Science 359, 689–692 (2018).

65. Suzuki, R. & Kawahara, H. UBQLN4 recognizes mislocalized transmembrane domain proteins and targets these to proteasomal degradation. EMBO Rep 17, 842–857 (2016).

66. Hessa, T. et al. Protein targeting and degradation are coupled for elimination of mislocalized proteins. Nature 475, 394–397 (2011).

67. Kriegenburg, F., Ellgaard, L. & Hartmann-Petersen, R. Molecular chaperones in targeting misfolded proteins for ubiquitin-dependent degradation. FEBS J 279, 532–542 (2012).

68. Abildgaard, A. B. et al. Co-Chaperones in Targeting and Delivery of Misfolded Proteins to the 26S Proteasome. Biomolecules 10, (2020).

69. McDonough, H. & Patterson, C. CHIP: a link between the chaperone and proteasome systems. Cell Stress Chaperones 8, 303–308 (2003).

70. Hilgarth, R. S. & Lanigan, T. M. Optimization of overlap extension PCR for efficient transgene construction. MethodsX 7, 100759 (2020).

71. Langmead, B. & Salzberg, S. L. Fast gapped-read alignment with Bowtie 2. Nat Methods 9, 357–359 (2012).

72. Krogh, A., Larsson, B., von Heijne, G. & Sonnhammer, E. L. Predicting transmembrane protein topology with a hidden Markov model: application to complete genomes. J Mol Biol 305, 567–580 (2001).

73. Tusnády, G. E. & Simon, I. Principles governing amino acid composition of integral membrane proteins: application to topology prediction. J Mol Biol 283, 489–506 (1998).

