## Supplementary Figures for "Conserved degronome features governing quality control associated proteolysis"

### Supplementary Figure 1

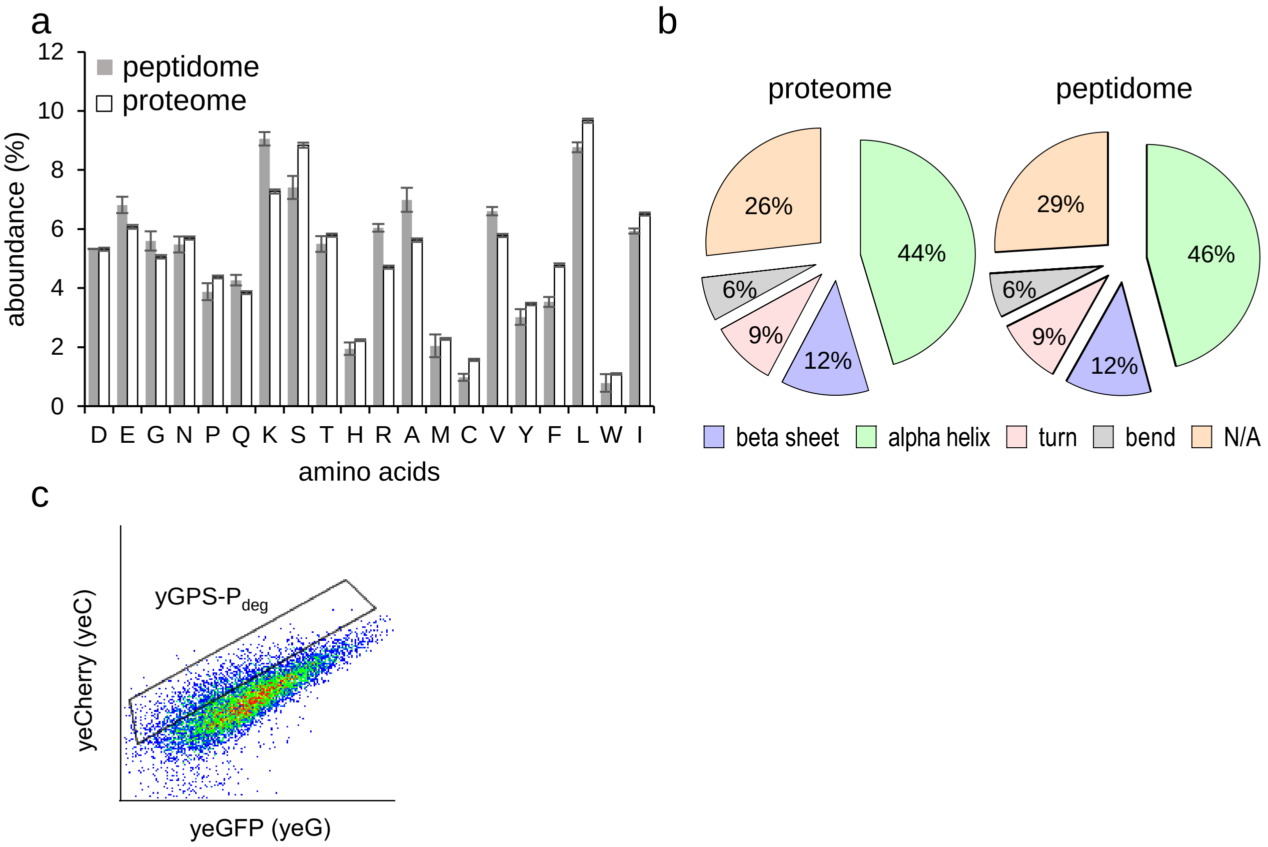

#### Supplementary Figure 2

Appears as a separate PDF file

### Supplementary Figure 3

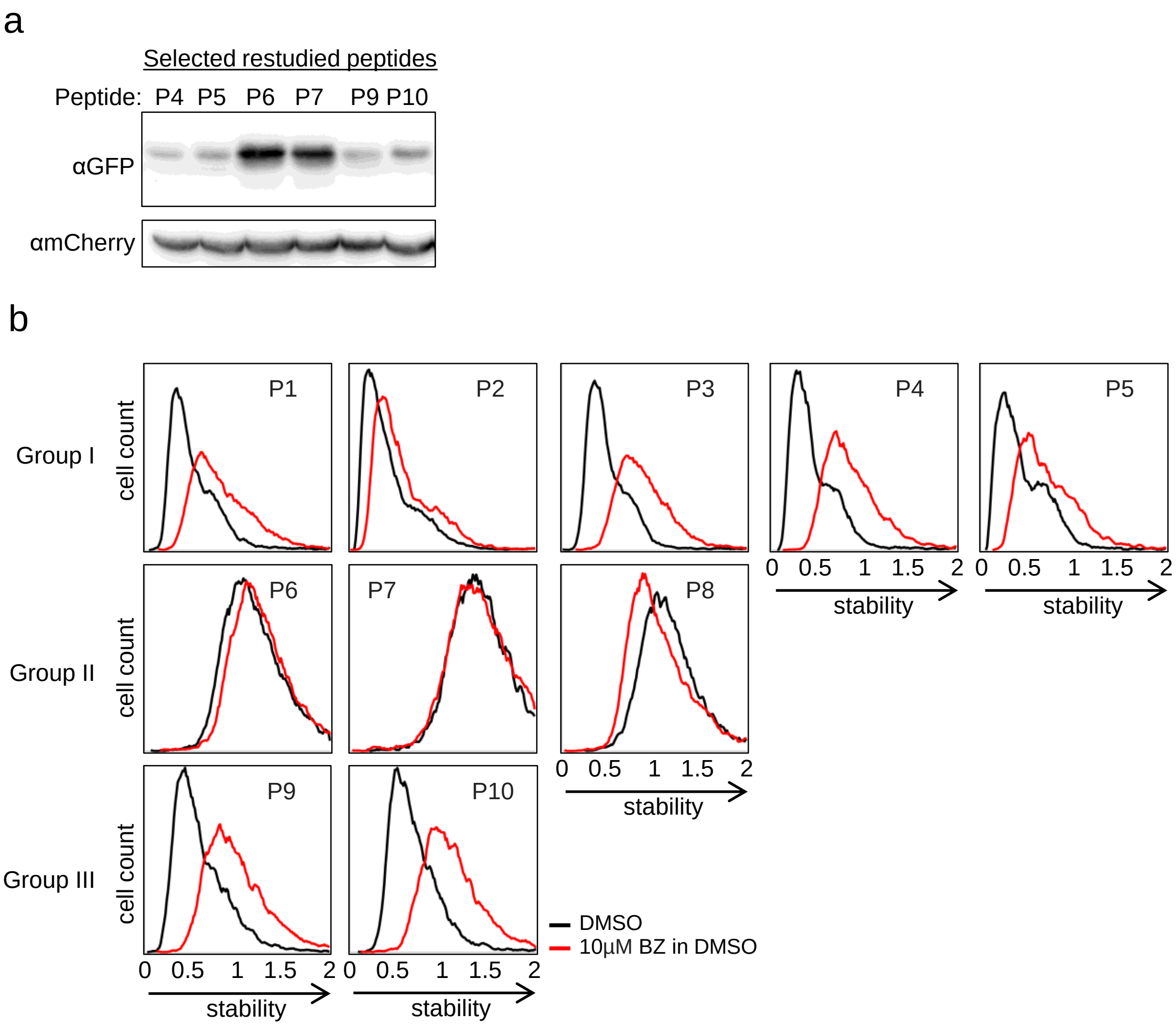

Supplementary Figure 4

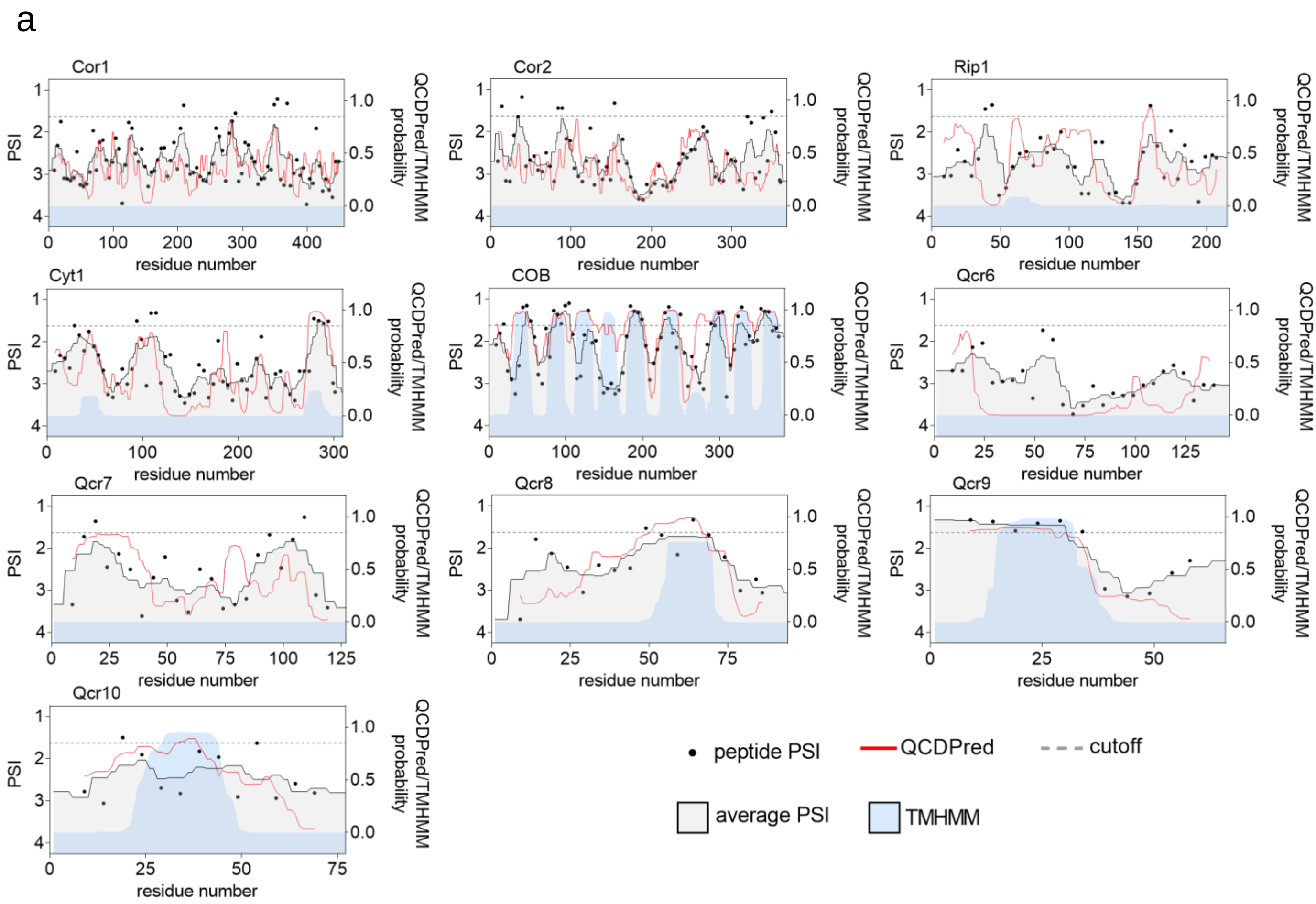


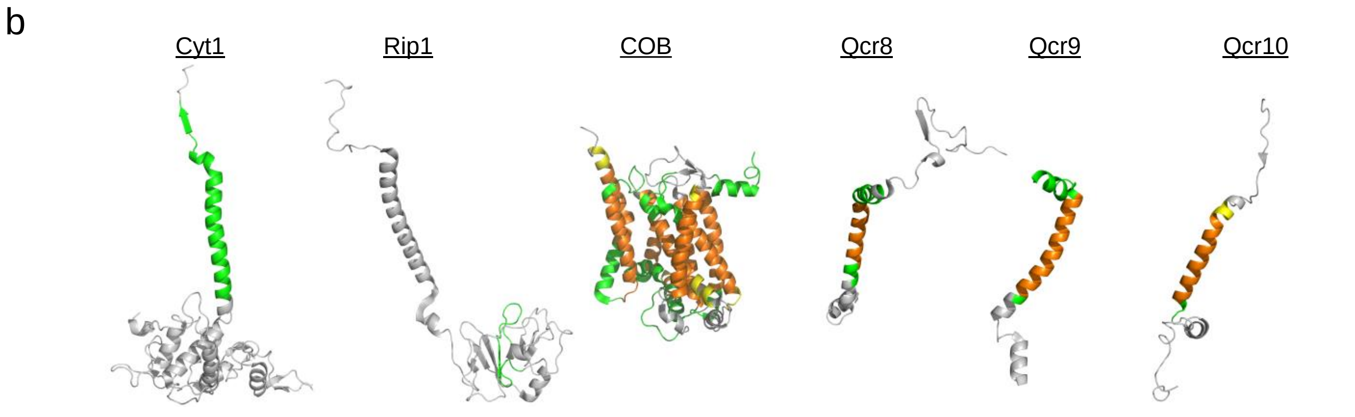

Supplementary Figure 5.

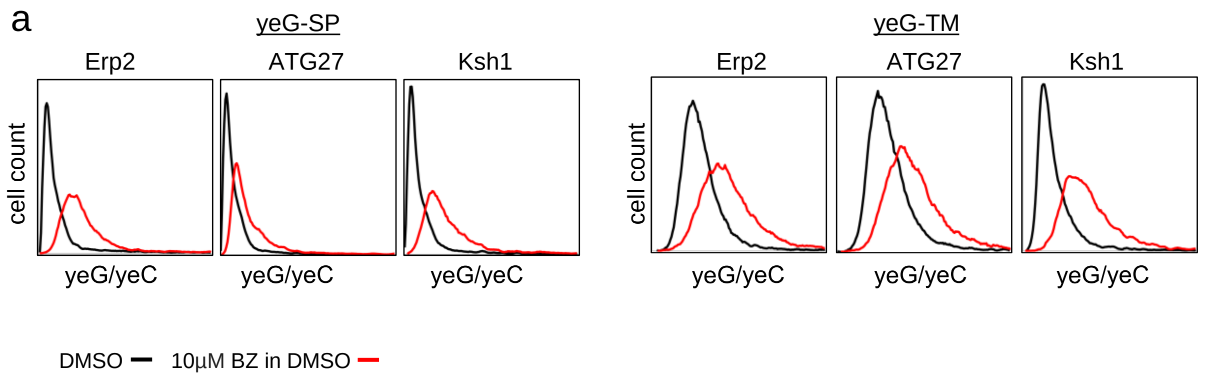


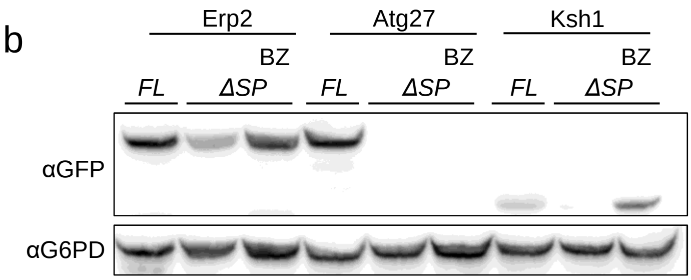


*Black*-DMSO
*Red*- 10µM BZ in DMSO

### Supplementary Figure 6

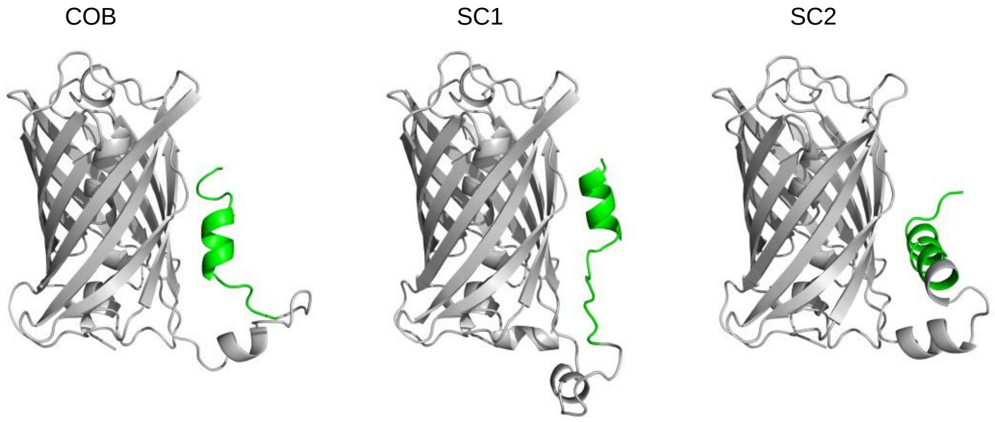

### Supplementary Figure 7

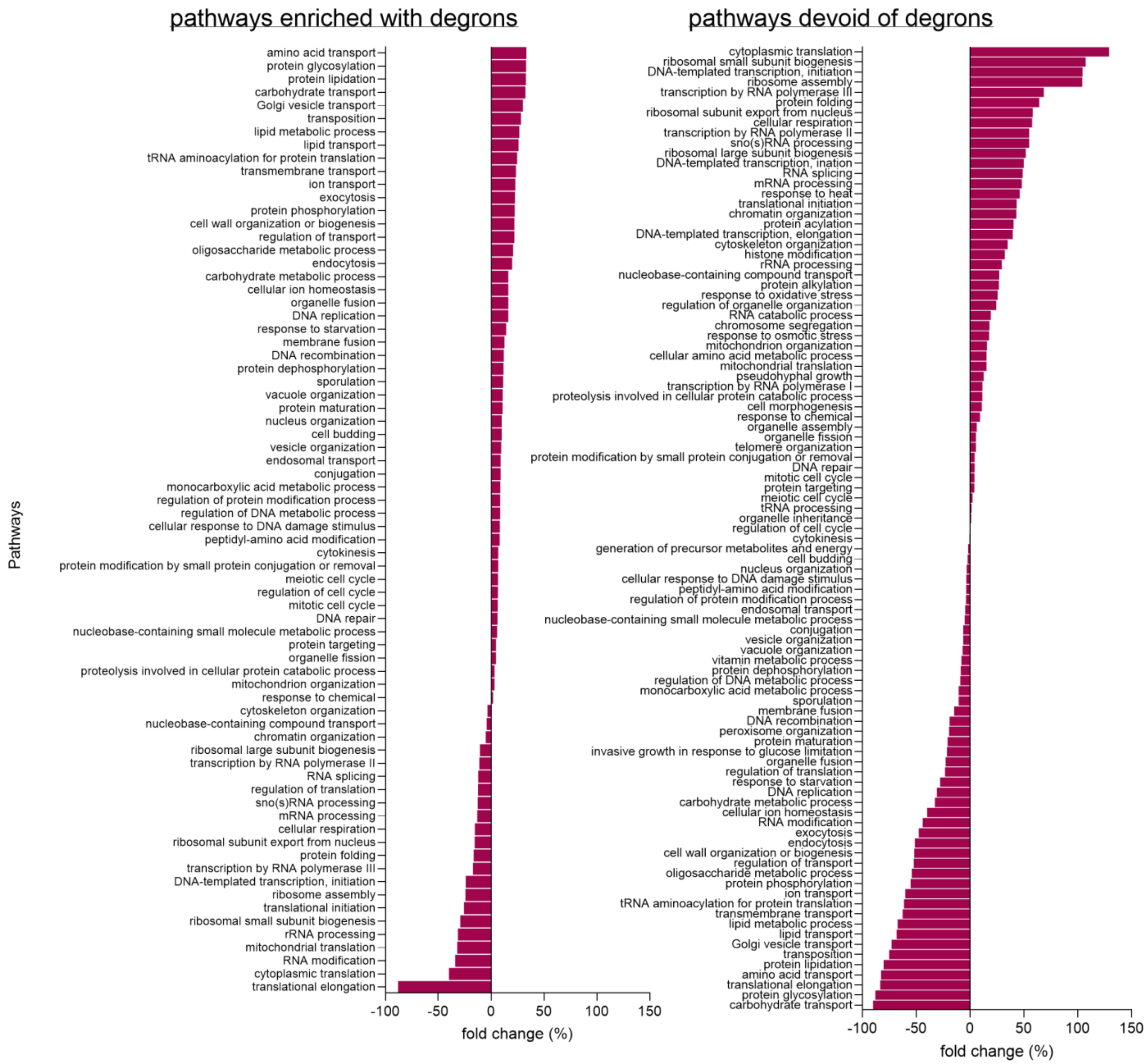

### Supplementary Figure 8

Cytosolic/nuclear Hsp40/70 chaperones

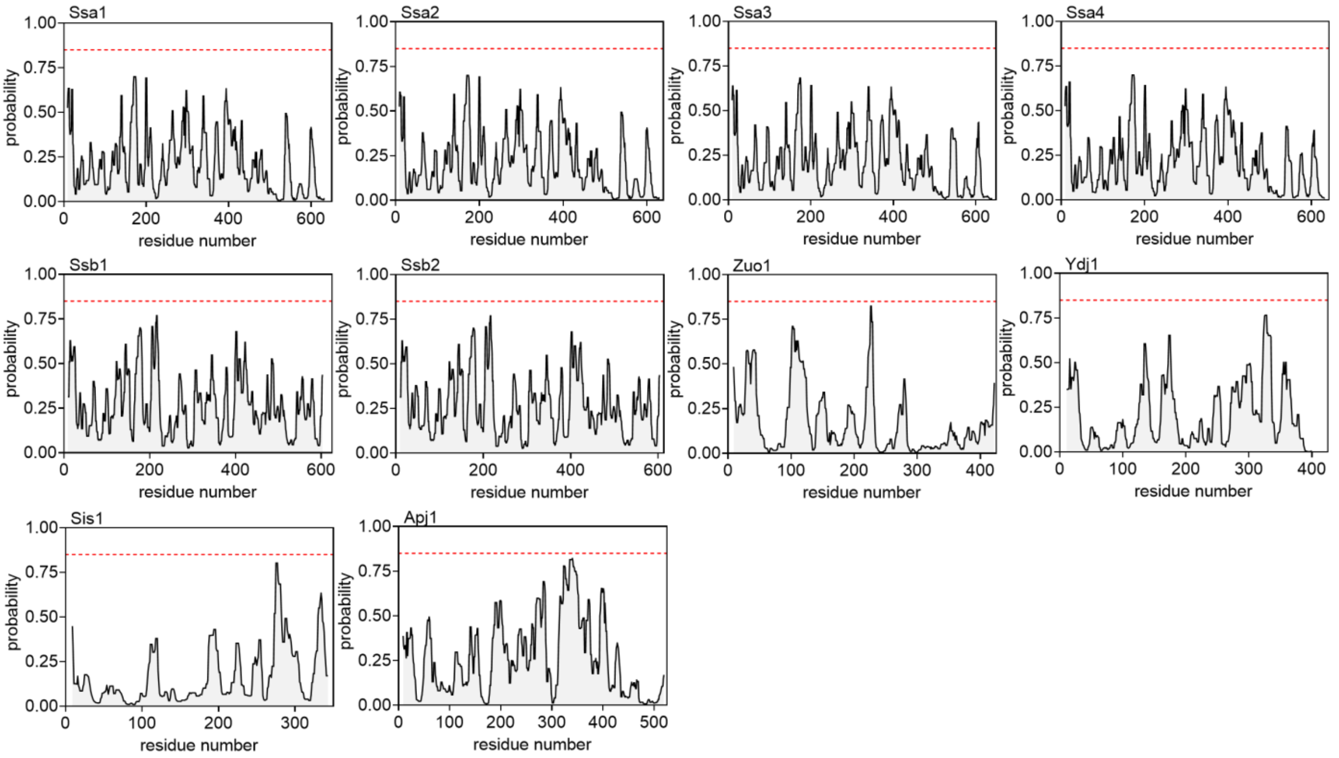


Cytosolic/nuclear Hsp90/100/110 chaperones

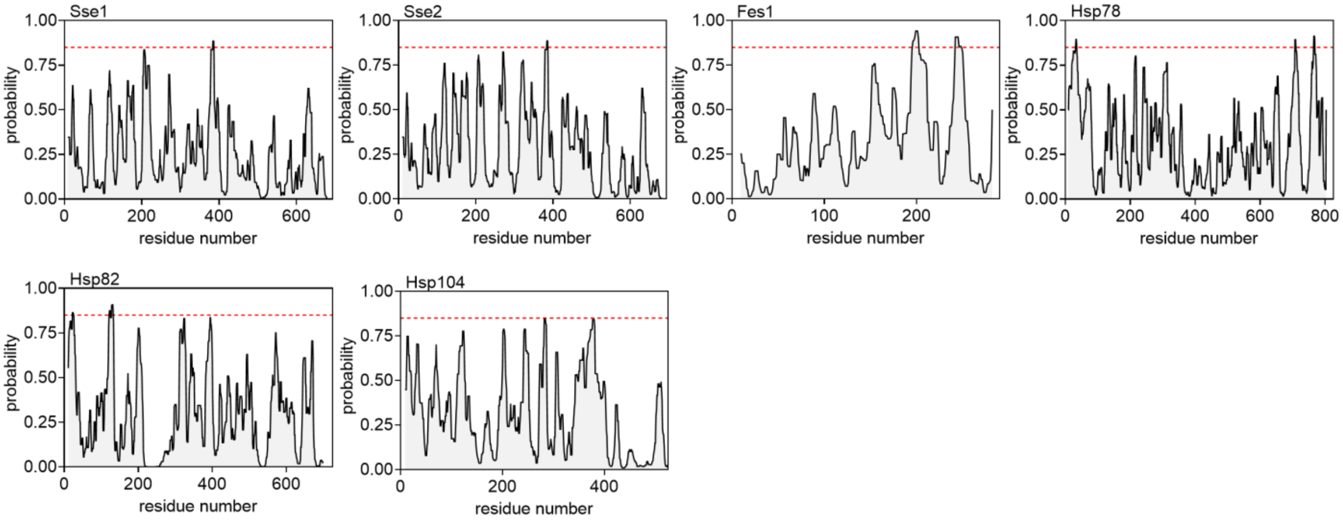

Supplementary Figure 9

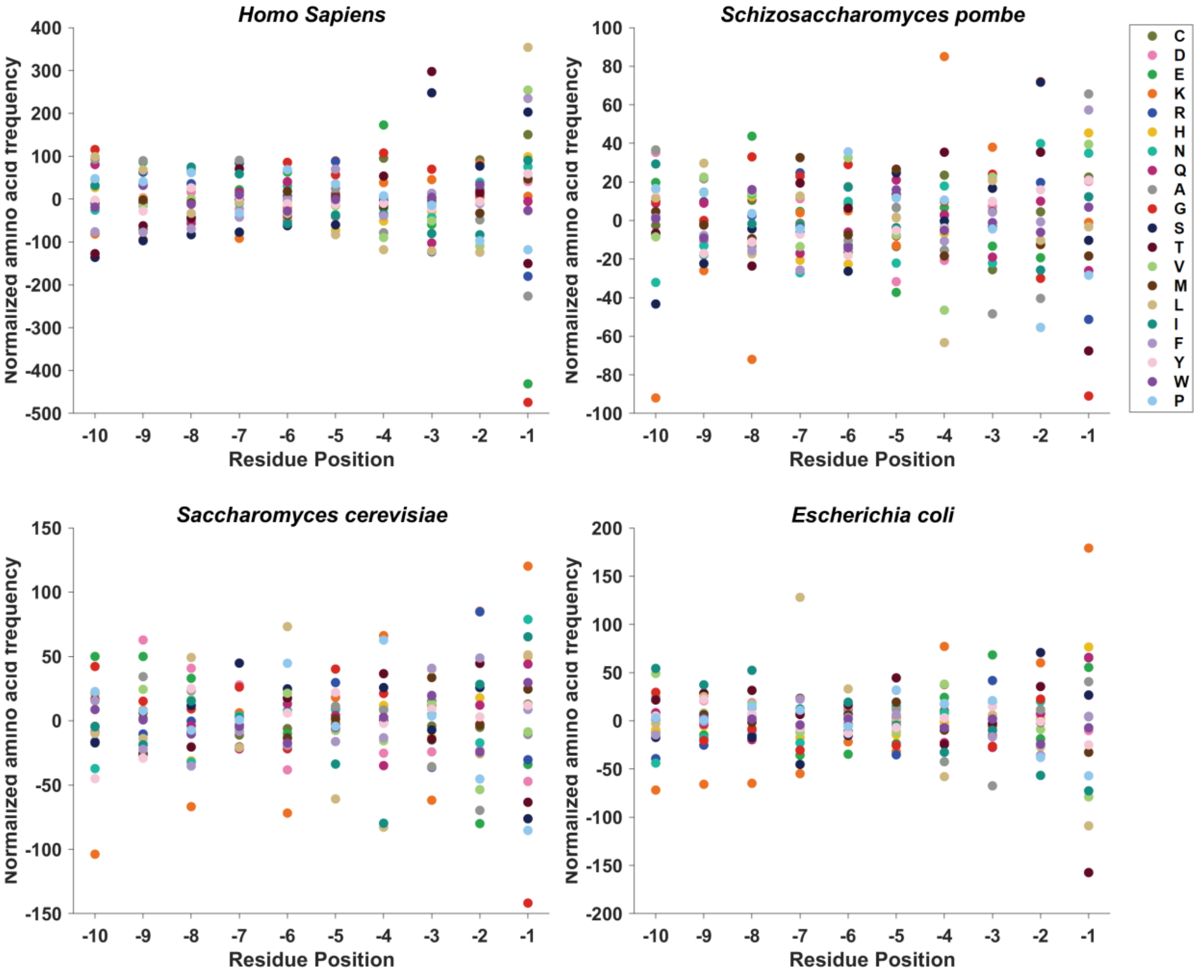
